# Bacterial defenses against a natural antibiotic promote collateral resilience to clinical antibiotics

**DOI:** 10.1101/2020.04.20.049437

**Authors:** Lucas A. Meirelles, Elena K. Perry, Megan Bergkessel, Dianne K. Newman

## Abstract

As antibiotic-resistant infections become increasingly prevalent worldwide, understanding the factors that lead to antimicrobial treatment failure is essential to optimizing the use of existing drugs. Opportunistic human pathogens in particular typically exhibit high levels of intrinsic antibiotic resistance and tolerance^1^, leading to chronic infections that can be nearly impossible to eradicate^2^. We asked whether the recalcitrance of these organisms to antibiotic treatment could be driven in part by their evolutionary history as environmental microbes, which frequently produce or encounter natural antibiotics^3,4^. Using the opportunistic pathogen *Pseudomonas aeruginosa* as a model, we demonstrate that the self-produced natural antibiotic pyocyanin (PYO) activates bacterial defenses that confer collateral tolerance to certain synthetic antibiotics, including in a clinically-relevant growth medium. Non-PYO-producing opportunistic pathogens isolated from lung infections similarly display increased antibiotic tolerance when they are co-cultured with PYO-producing *P. aeruginosa*. Furthermore, we show that beyond promoting bacterial survival in the presence of antibiotics, PYO can increase the apparent rate of mutation to antibiotic resistance by up to two orders of magnitude. Our work thus suggests that bacterial production of natural antibiotics in infections could play an important role in modulating not only the immediate efficacy of clinical antibiotics, but also the rate at which antibiotic resistance arises in multispecies bacterial communities.

## Introduction

The emergence and spread of bacterial resistance to clinical antibiotics is a growing public health concern worldwide^2^. Moreover, it is increasingly appreciated that antibiotic tolerance, defined as the ability to survive transient exposure to antibiotics, can also contribute to the failure of treatments for infections^5^. Yet bacterial resilience to antibiotics is anything but new: microbes in environments like soil have been producing natural antibiotics and evolving mechanisms of resistance and tolerance for millions of years^1,3^. We asked whether there could be a direct link between production of natural antibiotics by a human pathogen and recalcitrance of infections to clinical antibiotic treatment. One organism that fits this pattern is the opportunistic pathogen *Pseudomonas aeruginosa*, which is notorious for causing chronic lung infections in cystic fibrosis (CF) patients, as well as other types of infections in immunocompromised hosts^6^. *P. aeruginosa* produces several redox-active, heterocyclic compounds known as phenazines^7^. Despite possessing broad-spectrum antimicrobial activity^7^, including against *P. aeruginosa* itself^8^, phenazines have recently been shown to promote tolerance to clinical antibiotics under some circumstances, via mechanisms that have yet to be characterized^9,10^. Here, we sought to assess potential broader implications of the phenomenon by investigating whether phenazine-mediated tolerance to clinical antibiotics in *P. aeruginosa* is driven by cellular defenses that evolved to mitigate self-induced toxicity. In addition, we explored the ramifications of phenazine-induced tolerance for the evolution of bona fide antibiotic resistance, both in *P. aeruginosa* and in other clinically-relevant opportunistic pathogens.

## Results

### Responses to the self-produced natural antibiotic pyocyanin and their effects on tolerance to clinical antibiotics

We started by characterizing the defense mechanisms *P. aeruginosa* has evolved in response to its most toxic self-produced phenazine, pyocyanin (PYO)^7,8^. To do so in an unbiased fashion, we performed a genome-wide transposon sequencing (Tn-seq) screen in which the mutant library was exposed to PYO under starvation to maximize PYO toxicity^8^, and tolerance of the pooled mutants to PYO was assessed following re-growth (Fig. S1A). This revealed five broad categories of genes that significantly affect tolerance to PYO: (i) efflux systems, (ii) protein damage responses, (iii) membrane or cell wall biosynthesis, (iv) oxidative stress responses, and (v) carbon metabolism and transport (Fig. 1A). We validated the screen results by constructing and testing chromosomal clean deletion mutants for four of these genes (Fig. S1B). The strongest hits with a positive effect on PYO tolerance were transposon insertions in repressors of resistance-nodulation-division (RND) efflux systems (Fig. 1A, Fig. S1B, Table S1), which would cause overexpression of the downstream efflux pumps. Transposon insertions in the genes encoding the efflux pump proteins themselves did not have strong effects in our screen (Table S1), likely due to functional redundancy among the various efflux systems^11^. Indeed, when we challenged starved *P. aeruginosa* with PYO in the presence of the broad-spectrum RND efflux inhibitor phenylalanine-arginine β-naphthylamide (PAβN), cell death was dramatically accelerated, confirming that efflux pumps are necessary for minimizing PYO toxicity (Fig. 1B).

**Fig. 1.**
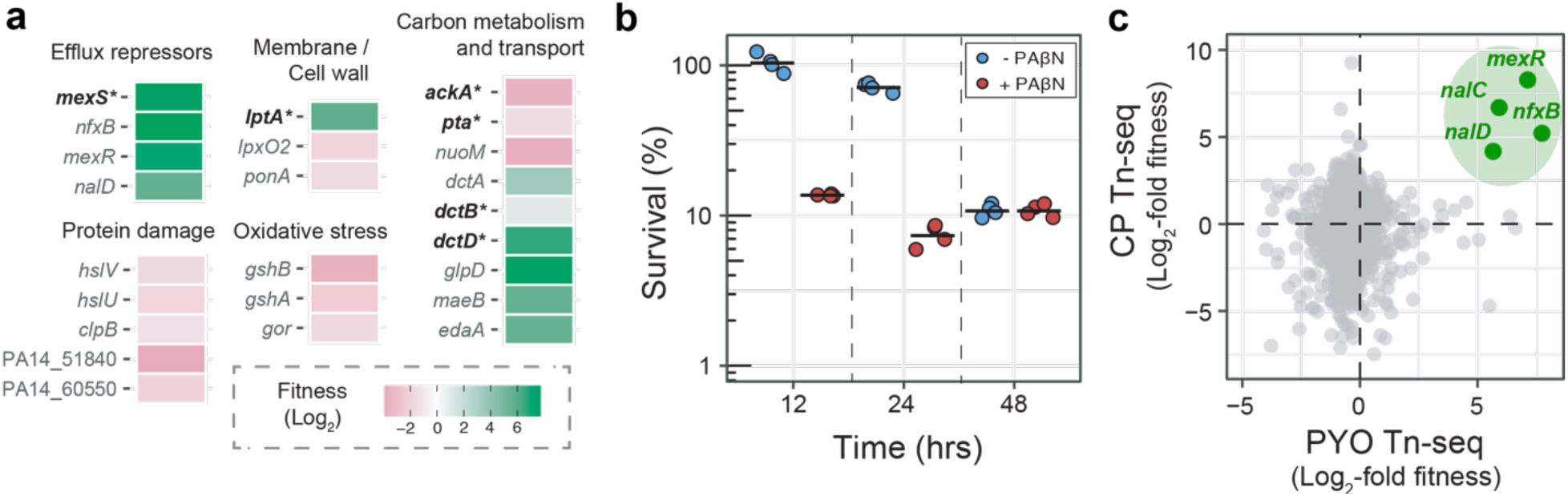
Mechanisms of tolerance to the self-produced natural antibiotic PYO in *P. aeruginosa*. **A**. Statistically significant fitness effects of transposon insertions in different representative genes under conditions that maximize PYO toxicity, as revealed by a genome-wide transposon sequencing (Tn-seq) screen (for experimental design, see Fig. S1A; for full dataset, see Table S1). See Methods for details on calculation of fitness. Asterisks show genes for which chromosomal clean deletion mutants were constructed and validated (Fig. S1B). **B**. Tolerance to PYO toxicity in the presence and absence of the efflux inhibitor PAβN. Each data point represents an independent biological replicate (n = 4), and the horizontal black lines mark the mean survival for each condition and time point. **C**. Fitness correlation analysis between PYO tolerance Tn-seq (this study) and CP persistence Tn-seq^13^. Efflux repressors present in both datasets are highlighted in green; *mexS* was not present in the CP Tn-seq dataset, while *nalC* is not shown in panel A because it was not statistically significant in our PYO Tn-seq despite its large effect on fitness, likely due to an insufficient number of unique transposon insertions in that gene. For full analysis, see Fig. S1C and Table S1.

Mutations in the efflux system repressors identified in our Tn-seq screen are commonly found in clinical isolates that are resistant to fluoroquinolone antibiotics, as the efflux systems regulated by these repressors efficiently export this class of drugs^12,13^. Likewise, these repressors were strong hits in a recent Tn-seq screen for genes that affect *P. aeruginosa* survival in the presence of the broad-spectrum fluoroquinolone ciprofloxacin^14^ (Fig. 1C, Fig. S1C, Table S1). We have also previously established that PYO upregulates expression of at least two efflux systems known to pump fluoroquinolones, *mexEF-oprN* and *mexGHI-opmD*^8,15^, in addition to the oxidative stress response genes *ahpB* (a thiol-specific peroxidase) and *katB* (a catalase)^8^; we confirmed the latter by performing qRT-PCR on the WT strain that produces PYO, a Δ*phz* mutant that does not produce PYO, and Δ*phz* treated with exogenous PYO (Fig. S2). Notably, phenazines and fluoroquinolones both contain at least one aromatic ring, unlike other antibiotics that are not thought to be pumped by *mexEF-oprN* and *mexGHI-opmD*, such as aminoglycosides^11^ (Fig. 2A). Thus, structural similarities could account for why efflux pumps that likely evolved to export natural antibiotics such as PYO can also transport certain classes of synthetic antibiotics. To determine whether PYO also induces other efflux systems known to pump clinical antibiotics besides fluoroquinolones, we performed qRT-PCR on representative genes from all 11 major RND efflux systems in the *P. aeruginosa* genome. These measurements confirmed that *mexEF-oprN* and *mexGHI-opmD* are the only two efflux systems significantly induced by PYO, and that the induction is PYO dose-dependent (Figs. 2B, S3, and S4). The *mexGHI-opmD* system in particular reached expression levels comparable to the constitutively-expressed *mexAB-oprM* efflux system (Figs. 2B-left and S3), which plays an important role in the intrinsic antibiotic tolerance and resistance of *P. aeruginosa*^11^.

**Fig. 2.**
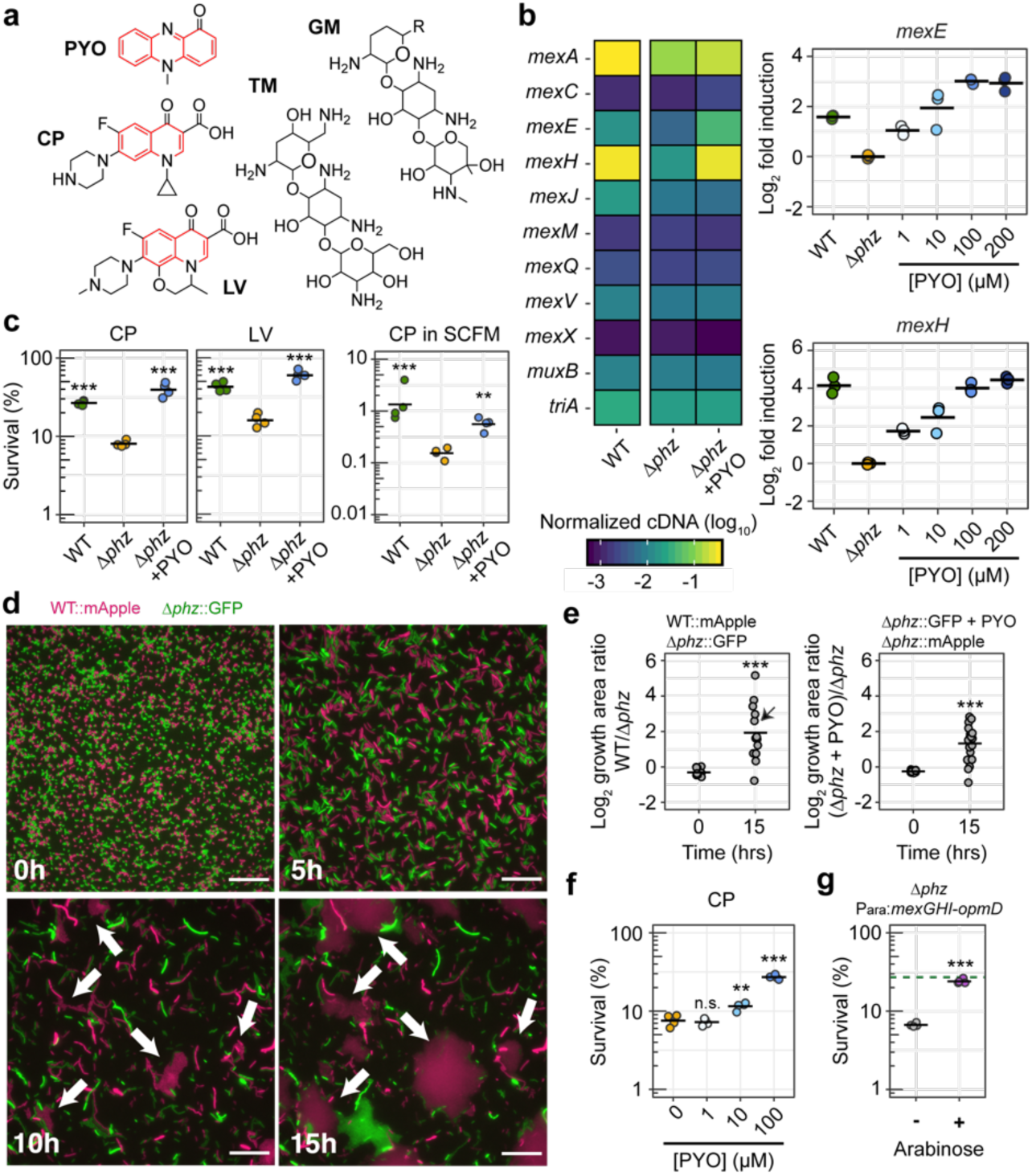
PYO induces expression of specific efflux systems, conferring cross-tolerance to fluoroquinolones. **A**. Structures of PYO, two representative fluoroquinolones (CP = ciprofloxacin, LV = levofloxacin) and two representative aminoglycosides (GM = gentamicin, TM = tobramycin). PYO and fluoroquinolones are pumped by MexEF-OprN and MexGHI-OpmD, while aminoglycosides are not^11^. Rings with an aromatic character are highlighted in red. **B**. Normalized cDNA levels for genes within operons coding for the 11 main RND efflux systems in *P. aeruginosa* (left). PYO-dose-dependent changes in expression of *mexEF-oprN* and *mexGHI- opmD* systems (right; n = 3). For full qRT-PCR dataset, see Figs. S2, S3 and S4. **C**. Effect of PYO on tolerance to CP and LV in glucose minimal medium (left), and to CP in SCFM (right) (all 1 µg/mL) (n = 4). PYO itself was not toxic under the experimental conditions^8^. WT made 50-80 µM PYO as measured by absorbance of the culture supernatant at 691 nm. See Fig. S5A for experimental design. **D-E**. Effect of PYO on lag during outgrowth after exposure to CP. A representative field of view over different time points (D; magenta = WT::mApple, green = Δ*phz*::GFP; see Movie S1) is shown together with the quantification of growth area on the agarose pads at time 0 hrs and 15 hrs (E). For these experiments, a culture of each strain tested was grown and exposed to CP (10 µg/mL) separately, then cells of both cultures were washed, mixed and placed together on a pad and imaged during outgrowth. The pads did not contain any PYO or CP (see Methods and Fig. S5D for details). White arrows in the displayed images point to regions with faster recovery of WT growth. The field of view displayed is marked with a black arrow in the quantification plot. The results for the experiment with swapped fluorescent proteins are shown in Fig. S5E. Scale bar: 20 µm. **F**. Tolerance of Δ*phz* to CP (1 µg/mL) in stationary phase in the presence of different concentrations of PYO (n = 4). **G**. Tolerance of Δ*phz* to CP (1 µg/mL) upon artificial induction of the *mexGHI-opmD* operon with arabinose (n = 4). The dashed green line marks the average survival of PYO-producing WT under similar conditions (without arabinose). Statistics: C, F – One-way ANOVA with Tukey’s HSD multiple-comparison test, with asterisks showing significant differences relative to untreated Δ*phz* (no PYO); E, G – Welch’s unpaired t- test (* p < 0.05, ** p < 0.01, *** p < 0.001, n.s. p > 0.05). In all panels with quantitative data, black horizontal lines mark the mean value for each condition. Individual data points represent independent biological replicates, except for in panel E, where the data points represent different fields of view.

Given that both efflux pumps and oxidative stress responses have previously been linked to antibiotic tolerance^16,17^, we asked whether the induction of these defenses by clinically-relevant concentrations of PYO^18^ could increase the tolerance of *P. aeruginosa* to clinical drugs. A survival assay (Fig. S5A) following treatment with clinical antibiotics revealed that, compared to the non- PYO-producing Δ*phz* mutant, the PYO-producing WT strain and PYO-treated Δ*phz* were more tolerant to both ciprofloxacin and another fluoroquinolone, levofloxacin (Fig. 2C). On the other hand, PYO did not confer increased tolerance to aminoglycosides (Fig. S5B), which are not substrates for the efflux pumps upregulated by PYO^11^. This finding contrasts with the conclusions of two previous studies on phenazine-mediated antibiotic tolerance, which claimed that phenazines broadly increase tolerance to all classes of antibiotics except cationic peptides^9,10^. However, one study focused on colony biofilms that produced only phenazine-1-carboxylic acid and phenazine- 1-carboxamide^9^, which are less toxic than PYO^8^ and consequently may induce a different set of cellular responses. Indeed, we found that aside from PYO, 1-hydroxyphenazine was the only other phenazine made by *P. aeruginosa* that increased tolerance to ciprofloxacin under our conditions, albeit to a lesser extent than PYO (Fig. S5C). The other previous study found that PYO increased planktonic culture cell densities in the presence of various antibiotics^10^, but these experiments did not directly demonstrate antibiotic tolerance^5^. Our results suggest that PYO preferentially induces tolerance to fluoroquinolones. Importantly, PYO also induced ciprofloxacin tolerance when *P. aeruginosa* was grown in synthetic cystic fibrosis sputum medium (SCFM) (Fig. 2C), suggesting that PYO production could contribute to antibiotic tolerance of this bacterium in CF patients.

Under *in vitro* conditions, PYO is typically produced in early stationary phase^15^. However, the heterogeneous nature of physiological conditions in infections^19,20^ could lead to intermixing of PYO-producing and -non-producing cells *in vivo*. We therefore tested whether exogenous PYO could increase the fluoroquinolone tolerance of cells harvested during log phase, which did not make PYO. To limit the growth of the no-antibiotic control, we exposed these cells to the antibiotics under nitrogen depletion. PYO still increased tolerance to both ciprofloxacin and levofloxacin under these conditions, suggesting that the induced tolerance phenotype does not depend on the metabolic state of stationary phase cells^8^ (Fig. S5B). Next, to visualize the recovery of cell growth after a transient exposure to ciprofloxacin, we performed a time-lapse microscopy assay (Fig. S5D). Interestingly, WT *P. aeruginosa* and PYO-treated Δ*phz* exhibited a shorter lag phase compared to non-PYO-treated Δ*phz* following ciprofloxacin treatment (Figs. 2D, E, S5E, and Movie S1), suggesting that PYO-induced defenses may help minimize cellular damage during the antibiotic treatment. We also found that addition of PYO to Δ*phz* increased ciprofloxacin tolerance in a dose-dependent manner (Fig. 2F), mirroring the dose-dependent induction of *mexEF-oprN* and *mexGHI-opmD* (Fig. 2A).

Given that PYO-induced efflux pumps transport specific substrates^11^, we asked if increased drug efflux could be the primary mechanism underlying PYO-mediated tolerance to fluoroquinolones. At high concentrations of ciprofloxacin, addition of the efflux inhibitor PAβN eliminated the survival advantage of PYO-treated cells, indicating that efflux pump activity is necessary for the PYO-mediated increase in antibiotic tolerance (Fig. S5F). Next, we constructed a Δ*phz* strain with the *mexGHI*-*opmD* operon under the control of an arabinose-inducible promoter (P_ara_:*mexGHI*-*opmD*). We verified that the transcription levels of *mexGHI-opmD* under arabinose induction were comparable to when PYO is present (Fig. S6). Indeed, arabinose induction of *mexGHI-opmD* expression increased ciprofloxacin tolerance to near-WT levels (Fig. 2G), suggesting that induction of this efflux system is sufficient to confer the PYO-mediated increase in tolerance. On the other hand, arabinose induction of the oxidative stress response genes *ahpB* or *katB* did not significantly increase tolerance of Δ*phz* to ciprofloxacin (Fig. S7). Importantly, the clinical relevance of *mexGHI-opmD* was previously not well known, as to our knowledge, there have been no reports of clinical mutants with constitutive overexpression of this efflux system. Taken together, our results demonstrate that PYO-mediated regulation of *mexGHI-opmD* expression modulates tolerance to clinically-used antibiotics in *P. aeruginosa*.

### Pyocyanin promotes the evolution of antibiotic resistance

Previous studies have demonstrated that mutations conferring antibiotic tolerance or persistence promote the evolution of antibiotic resistance^21,22^. Moreover, tolerance mutations can 1) interact synergistically with resistance mutations to increase bacterial survival during antibiotic treatment^23^ and 2) promote the establishment of resistance mutations during combination drug therapy^24^. Consequently, we performed fluctuation tests (Fig. 3A) to assess whether antibiotic tolerance induced by PYO could similarly promote the establishment of resistance mutations in populations of *P. aeruginosa* undergoing extended exposure to a clinical antibiotic. In clinical settings, antibiotic resistance is likely to result in treatment failure if a pathogen can grow at antibiotic concentrations above a threshold commonly referred to as a “breakpoint.” We adopted this criterion by selecting mutants on antibiotic concentrations either equal to or two-fold higher than the breakpoints defined by the European Committee on Antimicrobial Susceptibility Testing (EUCAST)^25^. Furthermore, we added PYO to our cultures either prior to the antibiotic selection step and/or concurrently with the antibiotic selection, in order to distinguish between the effects of preemptive versus continuous induction of PYO-regulated cellular defenses.

**Fig. 3.**
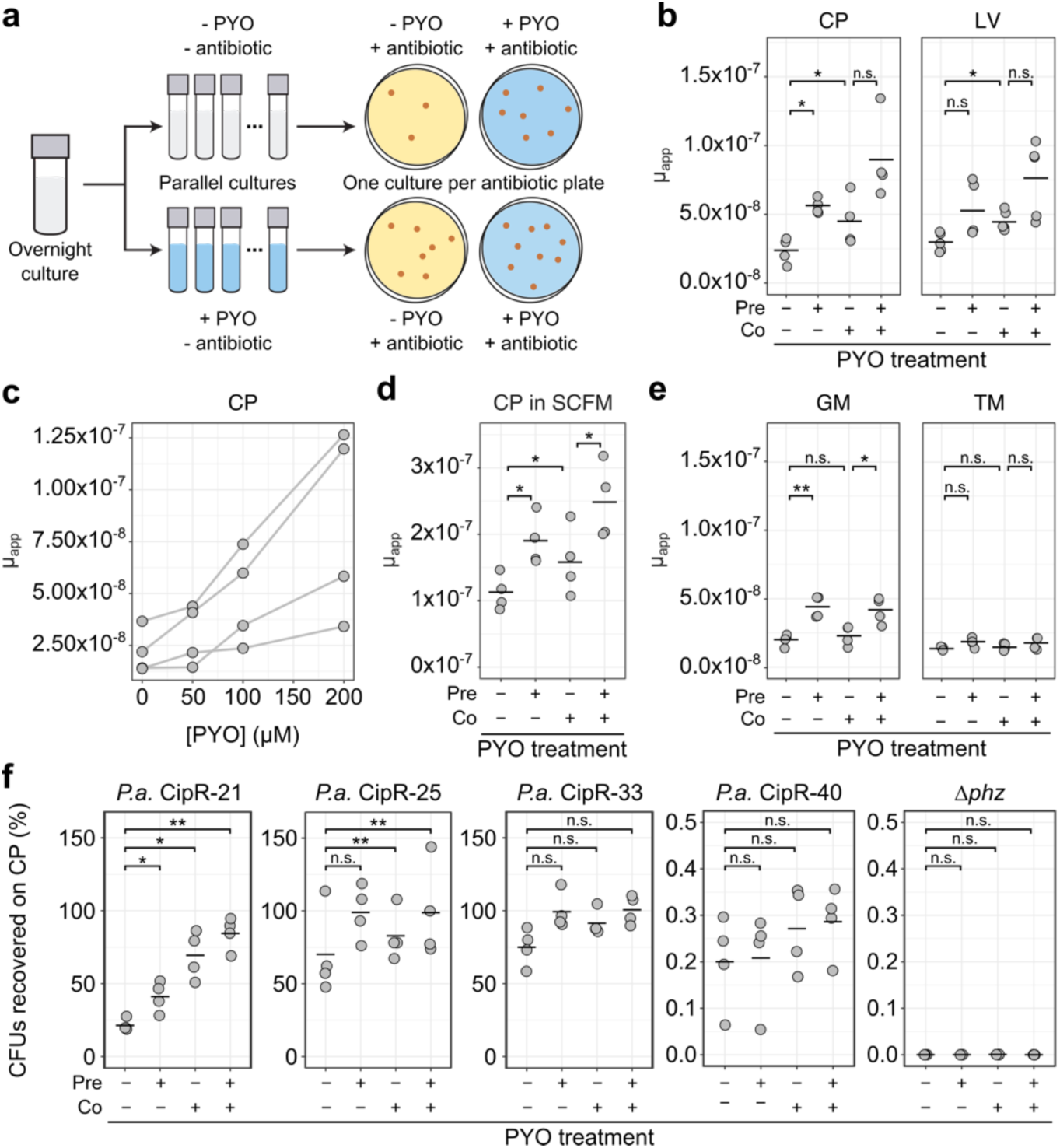
PYO increases the apparent rate of mutation to antibiotic resistance. **A**. Experimental design for fluctuation tests to determine the effect of PYO (100 µM unless otherwise noted) on apparent mutation rates. For panels B-E, mutation rates were calculated using an established maximum likelihood-based method that accounts for the effects of plating a small proportion of the total culture volume (see Methods for details). Each data point in those panels represents a single biological replicate comprising 44 parallel cultures. **B**. Apparent mutation rates of log-phase Δ*phz* grown in glucose minimal medium and plated on MH agar containing ciprofloxacin (CP, 0.5 µg/mL; n = 4) or levofloxacin (LV, 1 µg/mL; n = 5), with or without pre- and/or co-exposure to PYO relative to the antibiotic selection step. **C**. The apparent rate of mutation to resistance for Δ*phz* cells that were pre-treated with different concentrations of PYO and plated onto CP (0.5 µg/mL). Lines connect data points from the same biological replicate. **D**. Apparent mutation rates of log-phase Δ*phz* grown PYO in SCFM and plated on SCFM agar containing CP (1 µg/mL; n = 4) with or without pre- and/or co-exposure to PYO. **E**. Apparent mutation rates of log-phase Δ*phz* grown -/+ PYO in glucose minimal medium and plated during onto MH agar containing gentamicin (GM, 16 µg/mL; n = 4) or tobramycin (TM, 4 µg/mL; n = 4), with or without pre- and/or co-exposure to PYO. **F**. The percentage of CFUs recovered on CP (0.5 µg/mL) with or without PYO in the agar, for log-phase cultures of different partially-resistant mutants or Δ*phz* that were pre-grown with or without PYO in glucose minimal medium. Percentage recovery was calculated relative to total CFUs counted on non-selective plates. Data points represent independent biological cultures (n = 4). Statistics: B, D, E – Welch’s paired t-tests with Benjamini-Hochberg correction for controlling false discovery rate; F – Welch’s unpaired t-tests with Benjamini-Hochberg correction for controlling false discovery rate (* p < 0.05, ** p < 0.01, *** p < 0.001, n.s. p > 0.05). In B, C, E, and F, black horizontal lines mark the mean values for each condition.

Historically, mutation rates inferred from fluctuation tests have sometimes been assumed to correlate with the per-base mutation rate across the genome^26,27^. However, the results from these assays can also be affected by the number of unique possible mutations that permit growth under the selection condition^28^. To encompass both possibilities in this study, we use the term µ_app_ (apparent rate of mutation) as a proxy for the likelihood of evolving antibiotic resistance. We calculated this parameter using standard methods for fluctuation test analysis (see Methods for details). Regardless of whether PYO was added prior to or concurrently with the antibiotic selection, PYO increased µ_app_ for resistance to ciprofloxacin in both log-phase and stationary-phase cultures (Fig. 3B, Fig. S8A). These results indicate that pre-treatment with PYO was sufficient but not necessary to increase µ_app_ for fluoroquinolone resistance. Adding PYO at both stages of the fluctuation test resulted in an even greater increase in µ_app_ (Fig. 3B), while the increase in µ_app_ when PYO was added prior to antibiotic selection was dose-dependent (Fig. 3C). Similarly, cultures that were selected on levofloxacin also displayed an increased µ_app_ upon PYO treatment (Fig. 3B). More importantly, PYO significantly increased µ_app_ for cultures that were grown in liquid SCFM and selected on SCFM plates containing ciprofloxacin (Fig. 3D), suggesting that PYO produced by *P. aeruginosa* could promote mutation to antibiotic resistance in chronically infected lungs of CF patients^29^.

Because tolerance to aminoglycosides was not affected by PYO (Fig. S5B), we hypothesized that defense mechanisms induced by PYO would not promote mutation to aminoglycoside resistance. On the other hand, if PYO affected µ_app_ by acting as a mutagen, pre-treatment with PYO before antibiotic selection would be expected to increase µ_app_ for resistance to all classes of antibiotics. To differentiate between these modes of action, we repeated the fluctuation tests using gentamicin and tobramycin, representative members of the aminoglycoside family that disrupt protein translation^2^. Cultures that were pre-exposed to PYO exhibited increased µ_app_ for gentamicin resistance, although adding PYO directly to the antibiotic plates had no effect (Fig. 3E). However, neither pre-treatment with PYO nor co-exposure to PYO in the antibiotic plates significantly increased µ_app_ for tobramycin resistance (Fig. 3E). These differing responses to PYO depending on the choice of clinical antibiotic suggested that the observed changes in µ_app_ were related to the induction of cellular defenses rather than changes in the genome-wide per-base mutation rate. In fact, previous studies have suggested that gentamicin generates reactive oxygen species (ROS) more readily than tobramycin^30,31^. This could account for why pre-exposure to PYO increased µ_app_ for resistance to gentamicin but not tobramycin, given that PYO primes cells to detoxify ROS by inducing oxidative stress responses (Fig. S2). For resistance to fluoroquinolones, on the other hand, it is likely that simultaneous induction of multiple defenses is necessary to recapitulate the increases in µ_app_ upon exposure to PYO, as overexpression of individual genes induced by PYO did not significantly increase µ_app_ (Fig. S8B, C). Interestingly, this contrasted with our finding that induction of the *mexGHI-opmD* efflux system was sufficient to recapitulate PYO-mediated increases in fluoroquinolone tolerance. Together, these results suggest that resistance is shaped by a different set of cellular processes compared to tolerance, but that PYO-induced defense mechanisms nevertheless contribute to both types of resilience to antibiotic treatment.

We envisioned three ways in which, under antibiotic selection, PYO-induced defense mechanisms could lead to the apparent increases in mutation rates: A) by enhancing the growth of pre-existing partially-resistant mutants; B) by increasing the proportion of cells that survive and subsequently mutate to resistance; or C) by a combination of A and B. To distinguish between these scenarios, we implemented a two-pronged approach. First, to explore the possibility of scenario A, we searched for and isolated partially-resistant mutants from the fluctuation test plates containing ciprofloxacin. We re-grew these isolates both with and without PYO treatment and calculated the percentage of CFUs recovered on ciprofloxacin plates relative to non-selective plates, as a metric for each isolate’s level of resistance. Second, to determine the relative likelihoods of scenario A and scenario B, we examined the fit of our fluctuation test data to different formulations of the theoretical Luria-Delbrück (LD) distribution. Specifically, we compared mathematical models that make different assumptions regarding whether mutants arise prior to or during the antibiotic selection.

We identified several partially-resistant mutants for which the percentage of CFUs recovered on ciprofloxacin plates increased when the isolate was either pre-exposed or co-exposed to PYO (Fig. 3F). Whole genome sequencing revealed that these isolates contained mutations either in the efflux pump repressors *nfxB* or *mexS*, or in genes that affected growth rate, such as a ribosomal protein and a C4-dicarboxylate transporter (Table S3). As a validation of our assay, we also tested isolates with distinct colony morphologies that were not enriched following exposure to PYO in the original fluctuation tests. As expected, these mutants were fully resistant to ciprofloxacin at the original selection concentration (Fig. S8D). Sequencing these isolates again revealed mutations in *nfxB* or *mexS* (Table S3), albeit at different loci. Interestingly, PYO did not affect the ciprofloxacin resistance of at least one partially-resistant isolate, CipR-40, a slow-growth mutant containing a 19-bp deletion in a cell wall synthesis gene (Fig. 3F, Table S3). This suggests that PYO does not universally raise the baseline MIC of the entire population, but rather interacts synergistically with specific types of mutations conferring partial resistance. We repeated the stationary phase ciprofloxacin tolerance assay with these partially-resistant isolates and found that tolerance was also differentially affected by PYO (Fig. S8E). The tolerance and resistance phenotypes shared no underlying pattern, again indicating that tolerance is influenced by different cellular processes compared to resistance. Nevertheless, our results demonstrate that under antibiotic selection, a subset of partially-resistant mutants benefit from exposure to PYO.

That PYO increases µ_app_ by promoting the growth of pre-existing partially-resistant mutants was further supported by the alternative approach of evaluating the fit of our data to different mathematical models. Specifically, Pearson’s chi-square test indicated that our data closely fit the Hamon and Ycart model^32^ (Fig. S9, Table S2), which allows for differential fitness of mutants compared to WT cells, but assumes that all mutants arise pre-plating. However, we could not unequivocally rule out the possibility that post-plating mutations contributed to the increases in µ_app_, as a subset of our data also fit a mixed LD-Poisson model that assumes some mutations occurred during the antibiotic selection step^33^ (Fig. S9, Table S2). Together, these results suggest that the most probable explanation for PYO-mediated increases in apparent mutation rates is a combinatorial effect of increased detection of partially-resistant mutants and increased occurrence of post-plating mutations.

Importantly, even two-fold increases in mutation rates have been shown to significantly promote the evolution of resistance to high antibiotic concentrations (>10x original MIC)^34^. Moreover, it is well-established that partial resistance can rapidly lead to acquisition of full resistance via secondary mutations^35,36^. Indeed, several putative mutants appeared fully resistant to ciprofloxacin despite being enriched by exposure to PYO in the fluctuation tests (Fig. S8F). This discrepancy could be a result of acquiring secondary mutations either during growth on the original fluctuation test plates or during the pre-growth for the CFU-recovery assay. Thus, our results suggest that PYO may significantly affect the rate at which high-level resistance emerges in populations of *P. aeruginosa* undergoing long-term antibiotic exposure.

### Impacts of pyocyanin on antibiotic tolerance and resistance in other opportunistic pathogens

In both natural environments (e.g. soil) and clinical contexts (e.g. chronic infections), *P. aeruginosa* is found in multispecies microbial communities^37–40^. We hypothesized that microbes that frequently interact with *P. aeruginosa* would have evolved inducible defense mechanisms against PYO toxicity, and that production of PYO by *P. aeruginosa* might therefore also increase tolerance and resistance to clinical antibiotics in these community members. To test this hypothesis, we focused on the genera *Burkholderia* and *Stenotrophomonas*, both of which are (i) soil-borne gram-negative opportunistic pathogens that are frequently refractory to clinical antibiotic treatments^41–43^, and (ii) found in co-infections with *P. aeruginosa*, e.g. in CF patients^44^. Specifically, we tested a soil-derived strain, *Burkholderia cepacia* ATCC 25416; a non-CF clinical isolate of *Stenotrophomonas, S. maltophilia* ATCC 13637; and several clinical isolates of the three most prevalent *Burkholderia* species found in CF patients^37^: *B. cenocepacia, B. multivorans*, and *B. gladioli* (for descriptions of these strains, see Table S4).

We started by assessing each strain’s intrinsic resistance to PYO, as we expected that strong defenses against PYO toxicity would be required in order to benefit from exposure to this natural antibiotic. Indeed, for *S. maltophilia*, which was sensitive to PYO (Fig. S10A), the effects of PYO on antibiotic tolerance were complex: the presence of PYO was only beneficial when ciprofloxacin levels were low (1 µg/mL, which is below the MIC for this strain) (Fig. S10C). At a higher concentration of ciprofloxacin (10 µg/mL), PYO was detrimental in a dose-dependent manner (Fig. 4A), suggesting that the additional stress conferred by PYO outweighed any induction of defense mechanisms against ciprofloxacin. *S. maltophilia* also struggled to grow with *P. aeruginosa* in co-cultures (Fig. S10B, D), indicating that the conditions under which this species could potentially benefit from PYO are very limited. *B. cepacia, B. cenocepacia*, and *B. multivorans*, on the other hand, were highly resistant to PYO (Fig. S10A). For these three species, exogenously-added PYO increased tolerance to ciprofloxacin (Fig. 4B). Furthermore, for *B. cepacia*, we confirmed that this effect was PYO dose-dependent (Fig. 4B). We therefore tested whether *P. aeruginosa* could induce tolerance to ciprofloxacin in co-cultures with these *Burkholderia* strains. Using liquid culture plates in which the two species were separated by a permeable membrane (Fig. S10B), we found that PYO-producing *P. aeruginosa* strongly induced tolerance to ciprofloxacin in the *Burkholderia* species, and that the observed tolerance phenotypes were recapitulated by addition of exogenous PYO to co-cultures of *Burkholderia* and the *P. aeruginosa* Δ*phz* mutant, or to control cultures with *Burkholderia* alone in the same setup (Fig. 4C, Fig. S10E). Notably, for *B. cenocepacia* and *B. multivorans*, increased ciprofloxacin tolerance was also observed in co-cultures with a PYO-producing strain isolated from a CF patient, *P. aeruginosa* PA 76-11 (Fig. 4C, Fig. S10E). Moreover, similar results were obtained for co-cultures in SCFM (Fig. S10F). Together, these results suggest that PYO produced by *P. aeruginosa* in CF patients may decrease the efficacy of ciprofloxacin as a treatment for multispecies infections.

**Fig. 4.**
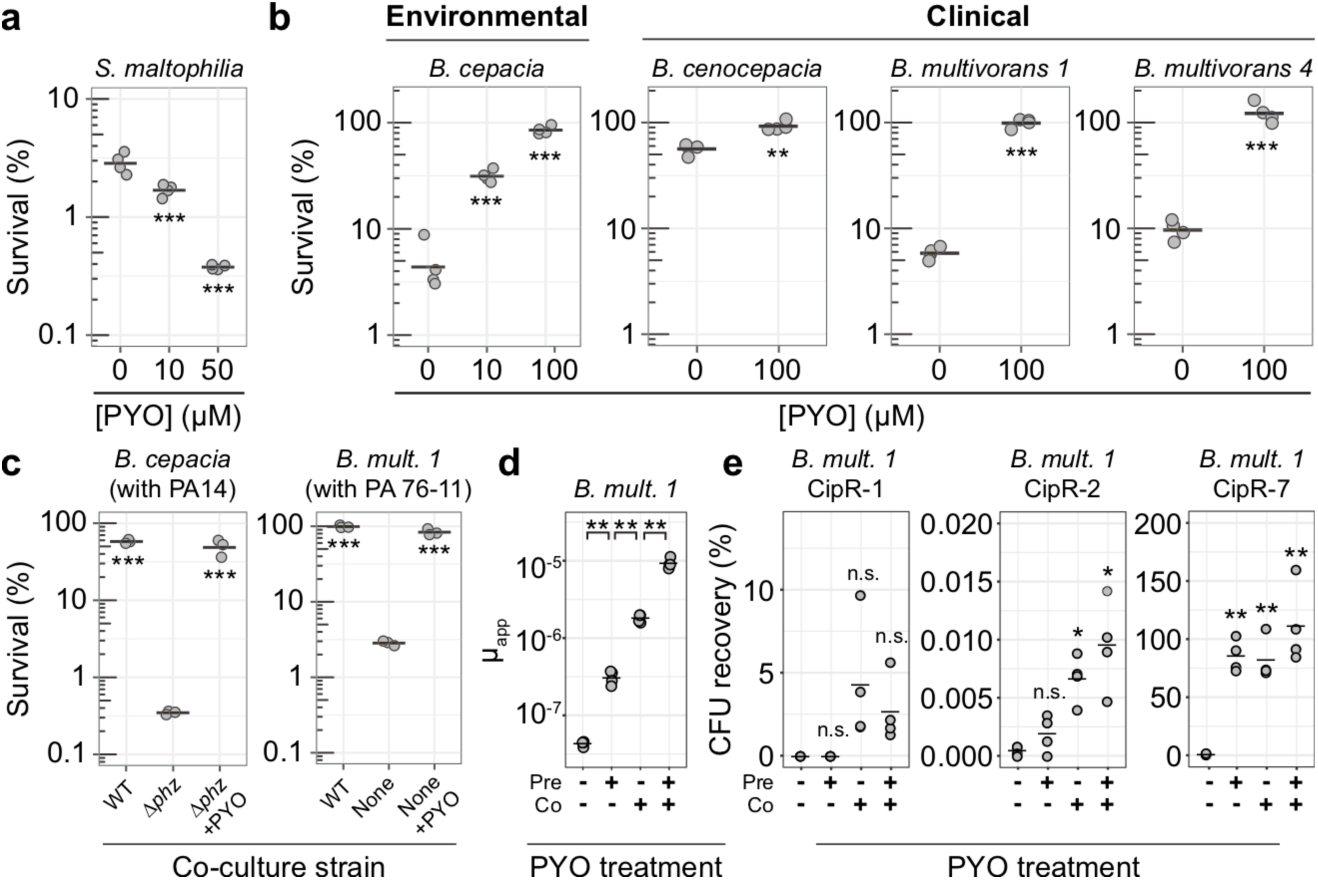
PYO promotes antibiotic tolerance and resistance in other opportunistic pathogens. **A**. Dose-dependent effect of PYO on *S. maltophilia* tolerance to high levels of ciprofloxacin (10 µg/mL; n = 4). **B**. Effect of PYO on the tolerance to ciprofloxacin (10 µg/mL) of multiple *Burkholderia* species isolated from environmental and clinical samples (n = 4). **C**. Effect of PYO produced by *P. aeruginosa* in co-cultures on the tolerance of different *Burkholderia* species to ciprofloxacin (10 µg/mL). PA14 is our model laboratory strain of *P. aeruginosa*, while PA 76-11 is a PYO-producing strain of *P. aeruginosa* isolated from a CF patient. The *Burkholderia* strains were plated separately for CFUs to assess survival following treatment with ciprofloxacin in the co-cultures (n = 3). See Fig. S10B for experimental design. **D**. The apparent rate of mutation to resistance when log-phase *B. multivorans 1* cells were plated on MH agar containing ciprofloxacin (8 µg/mL), with or without pre- and/or co-exposure to 100 µM PYO relative to the antibiotic selection step. Each data point represents a biological replicate comprising 44 parallel cultures (n = 4). **E**. The percentage of CFUs recovered on ciprofloxacin plates either with or without PYO in the agar, for exponential phase cultures of different partially-resistant *B. multivorans 1* mutants that were pre-grown with or without PYO in liquid cultures. Percentage recovery was calculated relative to total CFUs counted on non-selective plates (n = 4). Statistics: A, B, C – One-way ANOVA with Tukey’s HSD multiple-comparison test for comparisons of three conditions or Welch’s unpaired t-test for comparison of two conditions, with asterisks showing the statistical significance of comparisons with the untreated (no PYO) condition; D, E – Welch’s paired (D) or unpaired (E) t-tests with Benjamini-Hochberg correction for controlling false discovery rate (* *p* < 0.05, ** *p* < 0.01, *** *p* < 0.001). In all panels, data points represent independent biological replicates, and black horizontal bars mark the mean values for each condition.

We next asked whether PYO could mediate an increase in apparent mutation rate for ciprofloxacin resistance in *Burkholderia* species. We chose *B. multivorans* AU42096 (*B. multivorans* 1 in Fig. 4 and Fig. S10) as our model strain for these experiments because, among the clinical isolates, it displayed the strongest response to PYO in the ciprofloxacin tolerance assays. Remarkably, when selecting *B. multivorans* mutants on ciprofloxacin, we observed PYO-mediated increases in µ_app_ that were far more dramatic than for *P. aeruginosa*: pre-treatment with PYO increased µ_app_ for *B. multivorans* approximately 10-fold, while co-exposure to PYO in the antibiotic plate without pre-exposure increased µ_app_ approximately 40-fold, and the combination of pre- and co-exposure to PYO increased µ_app_ by 230-fold (Fig. 4D, Table S2). Notably, the magnitude of the latter effect is on par with observed differences between hypermutators, such as mutants deficient in the mismatch repair pathway, and their respective parent strains^45–47^. Moreover, hypermutators of *Burkholderia* isolated from CF infections are associated with clinical ciprofloxacin resistance^47^. In light of these observations, our results suggest that PYO could significantly affect clinical outcomes for co-infections of *P. aeruginosa* and *B. multivorans* treated with ciprofloxacin.

To verify that the *B. multivorans* colonies growing on ciprofloxacin in the presence of PYO were mutants, and to assess their responses to PYO, we isolated several putative mutants from the fluctuation test antibiotic plates and tested three in our CFU-recovery assay. All three displayed unique profiles of ciprofloxacin resistance in response to PYO treatment, as well as different maximal levels of resistance, though none were noticeably resistant to ciprofloxacin without exposure to PYO (Fig. 4E). Whole-genome sequencing of these isolates revealed that *B. multivorans* CipR-1 possessed mutations in three uncharacterized regulatory genes (Table S5). *B. multivorans* CipR-2 possessed mutations in two different homologs of the SpoT/RelA (p)ppGpp synthetase gene, which is known to affect antibiotic tolerance and resistance^48^. *B. multivorans* CipR-7 possessed a point mutation in DNA gyrase A (S83R), along with a point mutation in a malto-oligosyltrehalose synthase. Given that DNA gyrase A is the target of ciprofloxacin and that the specific mutated residue is likely homologous to the T83 residue that was mutated in a study of fluoroquinolone-resistant mutants in *B. cepacia*^49^, it is intriguing that this mutant was not fully resistant to ciprofloxacin in the absence of PYO; however, the specific amino acid substitution in this strain may have resulted in only a mild disruption of ciprofloxacin binding.

Lastly, we asked whether the *B. multivorans* mutants we detected primarily arose prior to or during the antibiotic selection. In all cases, the distribution of mutants closely matched the Hamon and Ycart formulation of the theoretical LD distribution, suggesting that the detected mutants arose prior to the antibiotic exposure (Fig. S11, Table S2). Interestingly, the Hamon and Ycart model also predicted the average relative fitness of mutants detected in PYO-treated samples to be significantly lower compared to mutants detected in non-PYO treated samples (Table S2; *p* < 0.05 for all three comparisons between non-PYO-treated and PYO-treated sample groups, using Welch’s paired t-test with Benjamini-Hochberg corrections for controlling the false discovery rate). In addition, unlike for *P. aeruginosa*, the mixed LD-Poisson distribution that allows for post-plating mutations was a poorer fit than the Hamon and Ycart model for all PYO-treated *B. multivorans* samples (Fig. S11, Table S2). Together, these results suggest that in *B. multivorans*, PYO increases µ_app_ by promoting growth of a wider range of mutants that arise prior to antibiotic selection, including those with slower growth rates.

## Discussion

Chronic infections are frequently caused by opportunistic pathogens whose long evolutionary history in the presence of natural antibiotics has potentially primed them for resilience against clinical antibiotic treatments. Indeed, many clinical antibiotic resistance genes are thought to have originated in environmental microorganisms as responses to microbial chemical warfare, with subsequent mobilization into human pathogens via horizontal gene transfer^1,3,4^. Here, we have demonstrated that tolerance and resistance to clinically-relevant concentrations of synthetic antibiotics can also arise as a collateral benefit of natural antibiotic production by an opportunistic pathogen.

*P. aeruginosa* is a particularly relevant example of an opportunistic pathogen whose self-produced natural antibiotics can promote tolerance and resistance to clinical antibiotics, given the large number of chronic infections caused by this bacterium worldwide^6^ and the fact that PYO has been detected in lung infection sputum samples at concentrations up to 130 µM^18^. Notably, treatments for infections caused by *P. aeruginosa* and other opportunistic pathogens often fail even when *in vitro* MIC tests indicate susceptibility to the chosen antibiotic^44^. Previous studies have attributed this discrepancy to metabolic and physiological changes within biofilms^50,51^, which represent a major form of bacterial life within infections^52^. Our results suggest that cellular defenses induced by bacterially-produced natural antibiotics may also contribute to *in vitro* versus *in vivo* differences in antibiotic susceptibility, as standard MIC tests are inoculated at a low cell density^25^, but *P. aeruginosa* typically does not make PYO *in vitro* until reaching a relatively high cell density^15^. Furthermore, the observation that PYO produced by *P. aeruginosa* strongly promotes antibiotic tolerance and resistance in *Burkholderia* species could hold important ramifications for the treatment of co-infections of these organisms in CF patients, for which clear best practices have yet to be established^44^. In particular, it could be prudent to avoid treating such infections with antibiotics for which PYO is likely to promote increased tolerance and resistance, such as fluoroquinolones, chloramphenicol, and trimethoprim/sulfamethoxazole—the latter two also being known substrates for efflux pumps that we have shown are upregulated by PYO^6,11^.

Finally, we note that our proposed model for collateral benefits of exposure to natural antibiotics (Fig. S12) potentially represents a broader phenomenon among human pathogens than has previously been appreciated. Many opportunistic pathogens originate in environments like soil^41,53^, where they have evolved in the presence of diverse natural antibiotics^1,3^, and *P. aeruginosa* is not the only pathogen with the capacity to synthesize its own antibiotics. For example, *Burkholderia* species possess the biosynthetic capability to produce a variety of compounds with antibacterial activity, whose potential clinical significance has not been explored^54^. If a given natural antibiotic induces efflux systems as a defense response, the only requirement for a consequent increase in tolerance to a clinically-relevant drug would be that the induced efflux system has some efficacy in transporting that drug—e.g. due to structural similarities like those shared by PYO and fluoroquinolones. This inference is supported by recent evidence that certain food additives or synthetic drugs antagonize the efficacy of specific clinical antibiotics by triggering stress responses in cells, including the induction of efflux pumps^55^. In fact, bacterially-produced toxic metabolites that promote antibiotic tolerance and resistance in human pathogens need not be limited to the types of molecules traditionally thought of as natural antibiotics. For example, indole secretion by highly antibiotic-resistant spontaneous mutants of *E. coli* enables partially-resistant mutants within the same species to grow at drug concentrations above their own MICs, in part by stimulating efflux pump expression^56^. Unlike PYO, indole is generally thought of as a signaling molecule rather than a natural antibiotic^57^, though it can likewise be toxic to bacteria at high concentrations^57,58^. Efforts to identify and characterize additional examples of such metabolites produced by opportunistic human pathogens could lead to an improved understanding of the modes of antibiotic treatment failure in clinics, and ultimately inform the design of more effective and longer-lived therapies.

## Acknowledgments

We thank members of the Newman lab and Shashank Gandhi for constructive feedback throughout the project and on the manuscript. We also thank Steven Wilbert for assistance with image analysis, David Basta for providing the plasmid used for *lptA* deletion, and The Millard and Muriel Jacobs Genetics and Genomics Laboratory at Caltech and Igor Antoshechkin for support during library preparation and sequencing of the Tn-seq samples. Finally, we thank John LiPuma (CFF *Burkholderia cepacia* Research Laboratory and Repository at the University of Michigan) for providing clinical *Burkholderia* strains. Grants to D.K.N. from the NIH (1R01AI127850-01A1) and ARO (W911NF-17-1-0024) supported this work. E.K.P. was supported by a National Science Foundation Graduate Research Fellowship under Grant No. DGE-1745301.

## Author contributions

These authors contributed equally and are listed alphabetically: Lucas A. Meirelles and Elena K. Perry. Study conception: L.A.M., E.K.P., D.K.N. Study design: L.A.M., E.K.P., M.B., D.K.N. Tn-seq: L.A.M, M.B. Tolerance experiments: L.A.M. Resistance experiments: E.K.P. Manuscript preparation: E.K.P., L.A.M., M.B., D.K.N. Study supervision and funding: D.K.N.

## Data and materials availability

Tn-seq data have been deposited at GEO under accession number GSE148769. Whole genome sequencing data for Δ*phz* and ciprofloxacin-resistant mutants of *P. aeruginosa* and *B. multivorans* AU42096 have been deposited at NCBI under accession number PRJNA625945. All other data that support the findings of this study are available from the corresponding author upon reasonable request. Correspondence and requests for materials should be addressed to.

## Methods

### Culture media and incubation conditions

Different culture media were used for different experiments as indicated throughout the Methods. Succinate minimal medium (SMM) composition was: 40 mM sodium succinate (or 20 mM, if specified), 50 mM KH_2_PO_4_/K_2_HPO_4_ (pH 7), 42.8 mM NaCl, 1 mM MgSO_4_, 9.35 mM NH_4_Cl, and a trace elements solution^59^. Glucose minimal medium (GMM) was identical to SMM, except with 10 or 20 mM glucose (as specified for different experiments) instead of succinate. SMM and GMM were prepared by autoclaving all components together for 20 min at 121°C, except for the carbon source and the 1000x trace elements stock solution, which were filter-sterilized and added separately; interestingly, we found that autoclaving MgSO_4_ with the other components was crucial for consistent PYO production by WT *Pseudomonas aeruginosa* UCBPP-PA14 in GMM. Luria-Bertani (LB) Miller broth (BD Biosciences) and BBL™Cation- Adjusted Mueller-Hinton II (MH) broth (BD Biosciences) were prepared according to the manufacturer’s instructions (notably with only a 10 min autoclave step for MH medium), with the addition of 1.5% Bacto™agar (BD Biosciences) to make solid media. Synthetic cystic fibrosis sputum medium (SCFM) composition was as described in previously^29^, with the addition of 3.6 µM FeSO_4_ and 0.3 mM N-acetyl-glucosamine^60^. All components except for the latter two were dissolved together, sterilized by filtration through a 0.22 µm membrane, and stored for up to two weeks; FeSO_4_ and N-acetyl-glucosamine solutions were prepared fresh each time or stored at -20 °C, respectively, and added to SCFM on the day of use. For SCFM agar, a 2x solution of the medium components was prepared and added to a separately autoclaved 3% molten agar solution, for a final concentration of 1x SCFM and 1.5% agar.

Antibiotics were prepared in concentrated stock solutions (100x or greater) and stored at - 20°C. Ciprofloxacin was dissolved in 0.1 M or 20 mM HCl, while levofloxacin, gentamicin, and tobramycin were dissolved in sterile deionized water. Phenylalanine-arginine β-naphthylamide (PAβN) dihydrochloride (MedChemExpress) was dissolved in sterile deionized water (50 mg/mL). Pyocyanin (PYO) was synthesized and purified as previously described^61,62^ and dissolved in 20 mM HCl to make 10 mM stock solutions. Experiments involving exogenous PYO always included negative controls to which an equivalent volume of 20 mM HCl was added. In addition, MH agar plates were buffered to pH 7 with 10 mM morpholinepropanesulfonic acid (MOPS) to avoid any pH changes upon addition of PYO or HCl; all other media used with exogenous PYO were already inherently buffered. Incubations were always done at 37°C, with shaking for liquid cultures (250 rpm), unless mentioned otherwise.

### Strain construction

In this study, we use PA14 as an abbreviation for UCBPP-PA14. *P. aeruginosa* PA14 was used for all experiments unless otherwise noted. For a full list of strains made in this study, see Table S4. Three types of strains were made in different *P. aeruginosa* PA14 backgrounds: (i) unmarked deletions, used for Tn-seq validation experiments; (ii) fluorescent strains for time-lapse microscopy experiments; and (iii) strains overexpressing one of the following three systems: *mexGHI-opmD, ahpB* and *katB*. Established protocols were used for all these procedures^63^.

Briefly, for unmarked deletions, ∼1kb fragments immediately upstream and downstream of the target locus were cloned using Gibson assembly into the pMQ30 suicide vector ^64,65^. Fragments amplified from *P. aeruginosa* PA14 genomic DNA (gDNA) and cleaned up using the Monarch PCR Purification kit (New England Biolabs) were used for Gibson assembly together with pMQ30 cut with SacI and HindIII. The assembled construct was then transformed into *Escherichia coli* DH10B, with transformants being selected in LB with 20 µg/mL gentamicin. All correctly-assembled plasmids were identified by colony PCR and verified by Sanger sequencing (Laragen). Next, for the insertion of the constructs into *P. aeruginosa* PA14 genome, tri-parental conjugation was performed following Choi and Schweizer^66^. All unmarked deletions were done in the *P. aeruginosa* PA14 Δ*phz* background (both *phzA-G1* and *phzA-G2* operons are deleted in this strain^15^), allowing clean experiments by addition of exogenous phenazines. Merodiploids containing the construct integrated into their genomes were selected on VBMM medium (3 g/L trisodium citrate, 2 g/L citric acid, 10 g/L K_2_HPO_4_, 3.5 g/L NaNH_4_PO_4_·4H_2_O, 1 mM MgSO_4_, 100 µM CaCL_2_, pH 7) with 100 µg/mL gentamicin following Choi and Schweizer^66^. Finally, merodiploids were then plated on LB lacking NaCl and containing 10% sucrose to select for colonies resulting from homologous recombination. Colonies missing the target locus (unmarked deletions) were identified by PCR. For all primers used, see Table S4.

Fluorescent strains used in time-lapse microscopy were made using previously published plasmids^63,67^. Constructs containing GFP and mApple florescent proteins under the control of the ribosomal *rpsG* gene were inserted in the *att*Tn7 site of *P. aeruginosa* PA14 Δ*phz* chromosome by tetra-parental conjugation, followed with selection on VBMM with 100 µg/mL gentamicin^66^.

Finally, overexpressing strains were made as previously described^63^. The previously made overexpression construct (pUC18T-miniTn7T-GmR vector containing the arabinose-inducible promoter P _ara_^63^) and the three different targets (*mexGHI-opmD, ahpB* and *katB*) were all amplified by PCR. Next, using Gibson assembly, the targets were cloned downstream of P_ara_ in the pUC18T-miniTn7T-GmR vector, resulting in the three different overexpression constructs: P_ara_:*mexGHI-opmD*, P_ara_:*ahpB*, and P_ara_:*katB*. The final constructs were introduced into the *att*Tn7 of the *P. aeruginosa* PA14 Δ*phz* background strain by tetraparental conjugation^66^.

### Transposon-sequencing (Tn-seq) experiment

The Tn-seq experiment was performed following the design presented in Fig. S1A. Two aliquots of the *P. aeruginosa* PA14 transposon library previously prepared^67^ were thawed on ice for 15 min, diluted to a starting optical density (OD_500_) of 0.05 in 50 mL of SMM, and grown aerobically under shaking conditions (250 rpm) at 37 °C for ∼ 4-5 generations to an OD_500_ of 0.8-1. These growing conditions were used for all the stages of the experiment. After growth in SMM, each aliquot was considered an independent replicate. Cells from each replicate were pelleted, washed and resuspended (5 mL in 18 × 150 mm glass tubes, OD_500_ = 2) in minimal phosphate buffer (MPB - 50 mM KH_2_PO_4_/ K_2_HPO_4_[pH 7], 42.8 mM NaCl) with and without 100 µM PYO. Cells were then incubated for 26 hrs under shaking conditions at 37° C. Therefore, the experiment consisted of four different samples that were later sequenced: (i) “R1 No PYO”, (ii) “R1 + PYO”, (iii) “R2 No PYO” and (iv) “R2 + PYO”. After the incubation, cultures from all treatments were pelleted, washed again to remove PYO, and resuspended in fresh SMM. Immediately, an aliquot of each sample was diluted to a starting OD_500_ of ∼ 0.05 in 25 mL SMM, followed by outgrowth for ∼ 4-5 generations to an OD_500_ of 0.8-1. After outgrowth, 2.5 mL of each sample was pelleted and stored at -80 °C.

Genomic DNA was extracted from the pelleted samples using the DNeasy Blood & Tissue kit (Qiagen). All the steps for sequencing library preparation followed exactly the protocol used by Basta et al.^67^, including (i) DNA shearing by sonication (to produce 200-500 bp fragments), (ii) end-repair, (iii) addition of poly(C) tail and (iv) enrichment of transposon-genome junctions and addition of adapter for Illumina sequencing by PCR^67,68^. The resulting amplified DNA samples were sequenced using 100 bp single-end reads on the Illumina HiSeq 2500 platform at the Millard and Muriel Jacobs Genetics and Genomics Laboratory at Caltech. Data analysis also followed Basta et al.^67^. In summary, sequences were mapped to the *P. aeruginosa* UCBPP-PA14 genome sequence using Bowtie 2^69^ and were analyzed in MATLAB using the ARTIST Tn-Seq analysis pipeline^70^, with non-overlapping windows of 100 bp across the genome^67,70^. Using Mann-Whitney U statistical test, the total reads mapping for each gene in the “+PYO” samples were compared to the corresponding reads in the “No PYO” control for each replicate independently^67,70^. Next, the read ratio for each replicate was calculated within ARTIST for each gene and then log_2_-transformed. Finally, the *p*-values for both replicates were combined using the Fisher’s combined probability test as done in Basta et al.^67^, and the average of the log_2_-ratios of the two replicates are also shown. For the log_2_-ratios and *p*-values for all PA14 genes, see Table S1. For heatmaps shown in Fig. 1A, the average log_2_-ratios (fitness) for the selected genes were plotted using the *geom_tile()* function from the ggplot2 package in R^71,72^.

### Tn-seq datasets correlation analysis

To compare the results of this Tn-seq analysis with a previously published study^14^ analyzing fitness determinants for survival during ciprofloxacin treatment in the *P. aeruginosa* PAO1 strain background (Fig. 1C, S1C and Table S1), the data from that study’s supplemental Table S1 were used. The normalized average ratio of reads in the treated sample compared to reads in the input sample for each gene (geometric mean of 3 replicates) was log_2_-transformed for comparison to the Tn-seq data described above. The list of genes was filtered to include only genes for which ratios were reported in both our PYO Tn-seq experiment and the ciprofloxacin Tn-seq study, and for which there are clear orthologs in both strains (n = 4209 genes). Orthologs were determined using the “pseudomonas.com” database^73^.

### Tn-seq validation experiments

To validate the Tn-seq results (Fig. S1B), experiments were performed by comparing survival of four different mutants (Δ*phz*Δ*ackA*Δ*pta*, Δ*phz*Δ*lptA*, Δ*phz*Δ*mexS* and Δ*phz*Δ*dctBD*) to the survival of the Δ*phz* strain upon exposure to PYO. The experimental design was very similar to the one used in for the Tn-seq, with minor adaptations. An overnight culture (5 mL) of each strain was grown in SMM (40 mM succinate) from LB plates. Cells were washed and re- suspended at an OD_500_ of 0.1 (or 0.25 for Δ*phz*Δ*dctBD*) in the same medium to start the new cultures (5 mL), which were grown to OD_500_ ∼ 0.8-1, pelleted, washed, and re-suspended in the same minimal medium without succinate (no carbon source) at OD_500_ of 1. For each strain, the culture was split across 8-12 wells (150 µL cultures) in a 96-well plate, with 100 µM PYO added to half of the cultures. 70 µL of mineral oil was added to the top of the wells to prevent evaporation. Propidium iodide (PI) at 5 µM was also added to the cultures to monitor cell death^8^. The plate was then moved to a BioTek Synergy 4 plate reader and incubated under shaking conditions at 37 °C for 24 hrs. After incubation, cultures were serially diluted in buffer and plated for colony forming units (CFUs) on LB agar, and survival in the presence of PYO was compared to the no-PYO control. Plates were incubated at room temperature (RT) and CFUs were counted after 36-48 hrs. In this study, a stereoscope was always used to count the CFUs. Survival levels were calculated for each mutant (i.e. for each replicate, the % survival for “+PYO” was calculated based on CFUs for “No PYO”). Then, the survival levels for each mutant were normalized by the survival levels of the Δ*phz* parent strain (i.e. % survival for “+PYO” for each mutant was divided by the average % survival for “+PYO” of the Δ*phz* strain); these “fitness” values were log_2_ transformed for plotting.

### PYO survival with efflux inhibitor

Survival assays with efflux inhibition were performed to test the importance of efflux systems in *P. aeruginosa* for tolerance against PYO toxicity. From a Δ*phz* overnight culture pre-grown in SMM (20 mM succinate), a new 7 mL culture was started in fresh SMM at an OD_500_ of 0.05 and was incubated for around 10 hrs (enough to reach stationary phase). Cells were then pelleted, washed and re-suspended in MPB at an OD_500_ of 1 (10 mL of culture was prepared). The culture was then split into four different treatments: (i) no PYO, no PAβN; (ii) 100 µM PYO, no PAβN; (iii) no PYO, with PAβN (50 µg/mL) and (iv) 100 µM PYO, with PAβN. Each of the treatments were split across 12 wells containing 150 µL of culture + 70 µL of mineral oil in a 96-well plate. The plate was incubated at 37 °C under shaking conditions using a BioTek Synergy 4 plate reader. Samples were serially diluted in MPB and plated for CFUs on LB agar after 12, 24 and 48 hrs. Survival for treatments containing PYO were calculated based on the CFUs counted for the negative control without PYO (Fig. 1C). At each time point, four wells were sampled, with each well considered an independent replicate. The experiment was repeated twice with similar results.

### Antibiotic tolerance experiments using P. aeruginosa

#### Tolerance assay for WT, Δ*phz* and Δ*phz* + PYO

For most antibiotic tolerance assays (except for tolerance using cells harvested during log-phase, see below), the experimental design shown in Fig. S5A was followed. WT and Δ*phz* cells were grown from a plate into an overnight culture in GMM. The overnight cultures were grown in GMM with 20 mM glucose. Next, cells were pelleted, washed and re-suspended at an OD_500_ of 0.05 (four replicates) in GMM (10 mM glucose) and incubated for around 20 hrs, reaching stationary phase, in 7 mL cultures (18 × 150 mm glass tubes). Three treatments were prepared: WT, Δ*phz* (no PYO) and Δ*phz* + 100 µM PYO. Cultures were then split into a negative control (no antibiotic) or antibiotic treatment (2 mL of culture per each treatment, using plastic Falcon tubes, VWR^®^ Cat. No. 352059). After addition of the antibiotic from concentrated stocks, cultures were incubated for four hours, serially diluted in MPB and then plated for CFUs on LB agar. Ciprofloxacin and levofloxacin were used at the concentrations mentioned in figure legends. Plates were incubated at RT and CFUs were counted after 36-48 hrs. Plates were always checked again after seven days to count any late-arising CFUs.

The same protocol was followed for the experiment testing different concentrations of PYO (Fig. 2F) and for the experiment testing how PYO impacts tolerance of different *P. aeruginosa* ciprofloxacin partially-resistant mutants (CipR-21, 25, 33 and 40, Fig. S8E). For the experiment testing tolerance after exposure to different phenazines (Fig. S5E), all the phenazines were dissolved in a common solvent (DMSO), which was used as the negative control; these experiments were performed in a Δ*phz\** mutant lacking not only the *phzA-G1* and *phzA-G2* operons but also all phenazine modification genes, to prevent the transformation of phenazine 1- carboxylic acid (PCA) into the other phenazines. For experiments performed in synthetic cystic fibrosis sputum medium SCFM (Fig. 2C), the same experimental design was followed, with the exception that SCFM was used instead of GMM.

#### Tolerance assay for strains with arabinose-inducible constructs

For these experiments (Fig. 2G and S7B), the 20 hrs cultures of each strain (Δ*phz* P_ara_:*mexGHI-opmD*, Δ*phz* P_ara_:*ahpB* and Δ*phz* P_ara_:*katB*) were grown with and without 20 mM arabinose for induction of the controlled systems, and then exposed to ciprofloxacin the same way described above. To rule out any non-specific interference of the inducer, negative controls with and without 20 mM arabinose using the parent Δ*phz* strain (without the insertions in the *att*Tn7 site) were also done. Adding arabinose to the Δ*phz* strain did not impact tolerance levels (Fig. S7).

#### Tolerance assay with PAβN

Experiments using the efflux inhibitor PAβN (Fig. S5F) were also performed similarly to the way as described above. The only differences were that after the 20 hrs incubation and before the addition of the antibiotic, PAβN was added to the cultures at a final concentration of 50 µg/mL. Cultures were incubated for 15 min and then ciprofloxacin was added, followed by four-hour incubation. For these experiments, instead of plating cells directly on LB, 1 mL of culture of each replicate/treatment was pelleted (12500 rpm for 2 min), washed in MPB for removal of CP and PAβN, and only then serially diluted in MPB and plated on LB for CFU counting.

#### Tolerance assay for cells harvested during log-phase

Δ*phz* cells were grown in overnight cultures in GMM (20 mM glucose). Next, cells were pelleted, washed and re-suspended into two new cultures, one with PYO (100 µM) and one without PYO, at an OD_500_ of 0.05 in GMM (10 mM glucose, 7 mL cultures). Cultures were grown until OD_500_ = 0.5 (around 5-6 hrs). Cells were then washed and re-suspended in the same medium at an OD_500_ of 0.5, but without the nitrogen source (i.e. no NH_4_Cl). PYO was re-added after washes to the culture that was pre-grown with PYO. The cultures, one with and one without PYO, were then split into different treatments: negative control (no antibiotic), ciprofloxacin (0.5 µg/mL), levofloxacin (1 µg/mL), gentamicin (16 µg/mL) and tobramycin (4 µg/mL). Then, they were all transferred to wells in a 96-well plate (three to four wells per treatment, with each well being considered an independent replicate). Cultures within wells contained 150 µL with an additional 70 µL of mineral oil on top to prevent evaporation. The depletion of nitrogen prevented growth in the negative control, which limited overestimation of the antibiotic killing effect (because survival rates were calculated relative to the negative control). The plates were incubated for four hours at 37 °C under shaking conditions (175 rpm) using a benchtop incubator (VWR^®^ incubator orbital shaker). The 96-well plate was kept inside an airtight plastic container with several wet paper towels to maintain high humidity attached to the shaker. After incubation, cells were serially diluted and plated on LB agar for CFU counting (Fig. S5B). A similar experiment was also performed with the strains containing arabinose-inducible constructs (Δ*phz* P_ara_:*mexGHI-opmD*, Δ*phz* P_ara_:*ahpB* and Δ*phz* P_ara_:*katB*) and the Δ*phz* background control (Fig. S7A), for which tolerance to ciprofloxacin (0.5 µg/mL) was tested. The experiment followed the same protocol described above, with the difference that, instead of presence or absence of PYO, strains were incubated in the presence or absence of 20 mM arabinose.

### Time-lapse microscopy experiment and quantification

Fluorescently tagged strains of WT or Δ*phz* were grown in GMM and tolerance experiments were performed as shown in Fig. S5D using ciprofloxacin (10 µg/mL). After the four-hour incubation with the antibiotic, cells were washed and re-suspended in GMM. The two different strains were then mixed and placed on an agarose pad containing GMM (no ciprofloxacin or PYO was added to the pad). Agarose pads were placed into a PELCO^®^ Clear Wall Glass Bottom Dish (Cat. No. 14023-20), and the dish was used for imaging within the microscope incubation chamber. Outgrowth was visualized using a Nikon Ti2E microscope with Perfect Focus System 4. Incubation proceeded for 12.5 to 15 hrs at 37 °C, with imaging every 15 min in bright field (phase contrast), green and red fluorescence channels (50 ms exposure with 470 nm LED lamp and a green-FITC filter [ex = 465-495nm, em = 515-555nm] for the GFP; 50 ms exposure with 555 nm LED lamp and a quad band filter [red ex = 543-566nm, red em = 580- 611nm] for mApple).

For image analysis, a Fiji macro was used (available upon request). Briefly, fluorescent channels (GFP/mApple) of the first and last time points were segmented using the “Auto Threshold” function and “Default” setting. The area of the segmented cells was then recorded using the “Analyze Particles” function in Fiji^74^. This allowed for quantification of the total area covered by cells within each channel, with each field of view being processed separately. After that, for each field of view, the total area covered by WT cells (or Δ*phz* + 100 µM PYO, depending on the experiment) was divided by the area covered by Δ*phz* cells to obtain the relative “growth area ratios”. This was done for first and last time points. Three experiments were performed, with different fluorescent protein/strain combinations: (i) WT::mApple/Δ*phz*::GFP (n = 13 fields of view, Fig. 2D, E, movie S1); (ii) Δ*phz*+PYO::GFP/Δ*phz*::mApple (n = 19, Fig. 2E); and (iii) WT::GFP/Δ*phz*::mApple (n = 16, Fig. S5E). GFP/mApple were controlled by the *rpsG* promoter for all of the strains (Table S4).

### RNA extraction and quantitative reverse transcriptase PCR (qRT-PCR)

#### Experiment 1 – measurement of PYO-induced gene expression

Six different treatments were prepared for this qRT-PCR experiment: (i) WT PA14, (ii) Δ*phz*, (iii) Δ*phz* + 1 µM PYO, (iv) Δ*phz* + 10 µM PYO, (v) Δ*phz* + 100 µM PYO and (vi) Δ*phz* + 200 µM PYO. Cultures of WT or Δ*phz* were grown overnight in GMM (20 mM glucose), then cells were washed and resuspended at an OD_500_ of 0.05 (three replicates) in fresh GMM (5 mL in culture tubes). Different concentrations of PYO were added to Δ*phz* cultures as mentioned and all cultures were incubated for around 8.5 hrs (until early stationary phase). This was enough time for WT to make PYO (around 50-70 µM, measured by absorbance at OD _691_^75^). After incubation, cells were pelleted, immediately frozen using liquid nitrogen and stored at -80 °C.

#### Experiment 2 – measuring arabinose induction of *mexGHI-opmD, ahpB* and *katB*

Eight different treatments were prepared for this qRT-PCR experiment, in which each of the four tested strains (Δ*phz*, Δ*phz* P_ara_:*mexGHI-opmD*, Δ*phz* P_ara_:*ahpB* and Δ*phz* P_ara_:*katB*) were incubated with and without 20 mM arabinose for artificial induction of the constructs. Cultures of the four strains were grown overnight in GMM (20 mM glucose), then cells were washed and resuspended at an OD_500_ of 0.05 (three replicates) in the same medium (5 mL in culture tubes), with and without 20 mM arabinose (for conditions without arabinose, the respective amount of water was added). Cultures were incubated for around 8.5 hrs, then pelleted, immediately frozen using liquid nitrogen and stored at – 80 °C.

For RNA extraction, previously published protocols were followed^8,63^. Briefly, samples were thawed in ice for 10 min and re-suspended in 215 µL of TE buffer (30 mM Tris.Cl, 1 mM EDTA, pH 8.0) containing 15 mg/mL of lysozyme + 15 µL of proteinase K solution (20 mg/mL, Qiagen), and then incubated for 8–10 min. For Lysis steps and RNA extractions it was used the RNeasy kit (Qiagen). Samples were then treated with TURBO DNA-free kit (Invitrogen) for removal of any contaminant gDNA. Next, cDNA was synthesized using iScript cDNA Synthesis kit (Bio-Rad) (1 µg of total RNA was used). For all these steps using kits, manufacturer’s instructions were followed. qRT-PCR reactions were performed using iTaq Universal SYBR Green Supermix (Bio-Rad) in 20 µL reactions using a 7500 Fast Real-Time PCR System machine (Applied Biosystems) following published protocols^8^. Standard curves for each primer pair were prepared using *P. aeruginosa* gDNA and were used for calculation of cDNA for each gene studied. The house-keeping gene *oprI* was used as a control gene for normalizations^63^.

Data showing total *oprI*-normalized cDNA levels (i.e. cDNA measured for a certain gene in a certain sample, divided by the respective cDNA measured for *oprI* in the same sample) and the log_2_-fold change in expression are shown in Figs. 2B and Figs. S2, S3 and S4. cDNA values for replicates within each efflux gene/treatment (shown in Fig. S3) were averaged and used with the *geom_tile()* function in R^71,72^ for generation of the heatmap shown in Fig. 2B.

Stenotrophomonas *and* Burkholderia *growth curves and antibiotic tolerance assays*

*Stenotrophomonas maltophilia* ATCC 13637, *Burkholderia cepacia* ATCC 25416, *B. cenocepacia* AU42085, *B. multivorans* AU42096 (*B. multivorans 1)*, and *B. gladioli* AU42104 were used in the growth experiments shown in Fig. S10A (for strain details, see Table S4). Each strain was grown overnight in GMM (20 mM glucose, 5 mL culture tubes) supplemented with 1x MEM amino acids (AA) (Sigma, Cat. No. M5550). Cells were pelleted, washed and re- suspended in new cultures at an OD_500_ of 0.05 in the same medium. Cultures were then split, different concentrations of PYO were added (0, 10, 50 or 100 µM for *S. maltophilia*; 0, 10 or 100 µM for all others), and moved to a 96-well plate (4 to 6 wells per treatment, with each well being considered an independent replicate). Cultures within wells contained 150 µL with an additional 70 µL of mineral oil on top to prevent evaporation. The plates were incubated at 37 °C under shaking conditions using a BioTek Synergy 4 plate reader with OD_500_ measurements every 15 min for 24 hrs to measure growth. Assays for tolerance to ciprofloxacin with or without exogenous PYO were performed for *S. maltophilia* (sensitive to PYO) and for four *Burkholderia* strains (all resistant to PYO): *B. cepacia, B. cenocepacia, B. multivorans 1* and *B. multivorans* AU18358 (*B. multivorans 4*). The experiments followed exactly what was done for *P. aeruginosa* (Fig. S5A), except that cultures were grown in GMM + AA, and are shown in Figs. 4A, 4B and S10C.

### Co-culture antibiotic tolerance experiments

To test how PYO produced by *P. aeruginosa* impacts tolerance to ciprofloxacin in other species, co-culture experiments were performed using membrane-separated 12-well tissue plate cultures containing 0.1 µm pore PET membranes (VWR^®^ Cat. No. 10769-226). Briefly, overnight cultures of the *P. aeruginosa* strain (WT/Δ*phz* PA14 or PA 76-11) and the respective other species tested (*S. maltophilia, B. cepacia, B. cenocepacia* or *B. multivorans 1*) were prepared in GMM (20 mM glucose) + AA. Cells were pelleted, washed and re-suspended to different ODs as follows: (i) for any *P. aeruginosa*-*Burkholderia* assay, *P. aeruginosa* starting OD_500_ = 0.05 and *Burkholderia* starting OD_500_ = 0.025; (ii) for the *P. aeruginosa*-*S. maltophilia* assay, *P. aeruginosa* starting OD_500_ = 0.01 and *S. maltophilia* starting OD_500_ = 0.1. *P. aeruginosa* was cultured in the bottom part of the well (600 µL), while the other species was cultured in the upper part of the well (100 µL), as shown in Fig. S10B. *B. cepacia* and *S. maltophilia* were cultured either with WT or Δ*phz P. aeruginosa* PA14 (with and without 100 µM PYO exogenously added). *B. cenocepacia* and *B. multivorans 1* were cultured either with PA 76-11 (a *P. aeruginosa* strain isolated from CF sputum that produced 50-100 µM PYO in these assays) or alone in the presence or absence of 100 µM PYO. For cases where *Burkholderia* was grown alone, the strain tested was grown in both the bottom and top parts of the membrane-separated wells. In all experiments, co-cultures were grown for around 20 hrs at 37 °C under shaking conditions (175 rpm) using a benchtop incubator, followed by addition of ciprofloxacin (concentrations were either 1 or 10 µg/mL, as specified in the figure legends) and incubation for four hours. The membrane-separated plates were kept inside an airtight plastic container with several wet paper towels to maintain high humidity attached to the shaker. For every co-culture combination in the membrane-separated plate, three wells were used as a negative control (no antibiotic) and three wells were used for ciprofloxacin treatment; each well was considered an independent replicate. After incubation with ciprofloxacin, cells were serially diluted in MPB and plated for CFUs on LB. In most cases, only *Burkholderia* cells were plated (Fig. 4C and Fig S10E). However, a co-culture experiment was also performed in SCFM (*P. aeruginosa* PA14 WT/Δ*phz* with *B. multivorans 1*) and in this case, both *P. aeruginosa* and *B. multivorans 1* were plated for CFUs (Fig. S10F). This experiment in SCFM followed the same overall experimental design used before, except for using SCFM instead of GMM in all steps.

### Determination of minimum inhibitory concentrations

MICs were determined using an agar dilution assay, in order to guide the choice of antibiotic concentrations for selecting *de novo* antibiotic-resistant mutants. Overnight cultures were grown for each strain in GMM (with 10 mM glucose) or GMM (10 mM glucose) + AA, respectively, then diluted to an OD_500_ of 0.5, from which 3 µL was spotted onto MH agar containing a 2-fold dilution series of the antibiotic. After the spots dried, the antibiotic plates were incubated upside-down for 48 hrs at 37°C before assessing the spots for growth. We considered the MIC to be the first concentration at which there was neither a lawn of background growth, nor dozens of overlapping colonies visible without magnification. We generally used 2x the MIC as the selection condition for fluctuation tests; for *P. aeruginosa*, this corresponded to the EUCAST resistance breakpoints for ciprofloxacin and levofloxacin, while our chosen concentrations of gentamicin and tobramycin were two-fold higher than the EUCAST breakpoints^25^. EUCAST breakpoints are not available for *Stenotrophomonas* spp. or the *Burkholderia cepacia* complex. The appropriateness of the selection condition for was additionally verified by performing a fluctuation test, as described below, and choosing the antibiotic concentration that reliably yielded a countable number of colonies (zero to several dozen, with at least several non-zero counts per 44 parallel cultures) in each well.

### Fluctuation tests, calculation of mutation rates, and model fitting

For all tested strains and conditions, fluctuation tests were performed by inoculating 200 µL cultures in parallel in a flat-bottomed 96-well plate. All reported fluctuation test data for *P. aeruginosa* are from experiments using the Δ*phz* strain. We also performed fluctuation tests using the *P. aeruginosa* PA14 WT strain, and performed phenotypic and genotypic characterization of partially-resistant mutants detected in those experiments (see below); however, the effect of PYO on apparent mutation rates in WT was difficult to interpret due to inconsistent PYO production in the 96-well plates. For cultures that were grown with PYO (or arabinose in the case of strains with arabinose-inducible constructs), the PYO (or 20 mM arabinose) was added to the medium before inoculation. The cultures were inoculated with a 10^−6^ dilution of a single overnight culture (representing a biological replicate) that had first been diluted to a standard OD_500_ of 1.0, corresponding to an initial cell density of approximately 2000- 2500 CFUs/mL (400-500 cells/culture). Each treatment condition consisted of 44 such parallel cultures. The 96-well plates were placed inside an airtight plastic container with several wet paper towels to maintain high humidity, then incubated at 37°C with shaking at 250 rpm. For plating during log-phase, the cultures were incubated until reaching approximately half-maximal density (OD_500_ of 0.4-0.7 for *P. aeruginosa* in GMM with 10 mM glucose, or 0.9-1.2 for *P. aeruginosa* in SCFM or *B. multivorans* in GMM + AA). For plating during stationary phase, the cultures were incubated for 24 hrs. The cultures were then plated by spotting 40-50 µL per culture into single wells of 24-well plates (for any given experiment, the same volume was spotted for all parallel cultures); each well contained 1 mL of MH agar or SFCM agar plus an antibiotic, with or without 100 µM PYO (or 20 mM arabinose for strains with arabinose-inducible constructs). In the case of *B. multivorans* cultures that were spotted onto antibiotic plates containing 100 µM PYO, the cultures were first diluted 1:10 (if not pre-treated with 100 µM PYO) or 1:100 (if pre-treated with 100 µM PYO). At the same time as plating onto the antibiotic plates, six representative cultures from each treatment were serially diluted and plated on LB agar plates to assess total CFUs. The antibiotic plates were incubated upside down, in stacks of no more than eight, at 37°C for 16-24 hrs for *P. aeruginosa* (except for gentamicin plates, which were incubated for 40-48 hrs) or 40-48 hrs for *B. multivorans*. Subsequently, colonies were counted under a stereoscope at the highest magnification for which the field of view still encompassed an entire well; occasionally, a well contained too many colonies to count (a so-called “jackpot” culture^76^), in which case that culture was discarded from further analysis. The LB agar plates for total CFU counts were incubated for 30-36 hrs at RT before counting colonies at the same magnification.

Mutation rates reported in the figures were calculated using the function newton.LD.plating from the R package rsalvador^77^ to estimate *m*, the expected number of mutations per culture. This is a maximum likelihood-based method for inferring mutation rates from fluctuation test colony counts, based on the classic Luria-Delbrück (LD) distribution with a correction to account for the effects of partial plating (i.e. plating a portion of each culture rather than the total volume)^77^. We chose this method because it has been shown to be the most accurate estimator of *m* when partial plating is involved^77,78^. To get µ_app_ (apparent mutation rate per generation) from *m*, we divided *m* by the total number of cells per parallel culture^77^, as estimated from the mean number of CFUs counted for the six representative cultures.

To compare the fits of different formulations of the LD distribution to our data, we generated theoretical cumulative distributions using the parameter values estimated for our data. Specifically, for the Hamon and Ycart version of the LD model^32^, we estimated *m* and *w* (relative fitness of mutants compared to the parent strain in the non-selective pre-plating liquid growth medium) using the function GF.est from the R script available at http://ljk.imag.fr/membres/Bernard.Ycart/LD/ (version 1.0; note that in the script, *m* is called alpha and 1/*w* is called rho); then, we used the function pLD from the same script to generate the theoretical distribution. For the mixed LD-Poisson and basic LD models, we wrote and used an R translation of the MATLAB code written by Lang *et al*.^33^; the original code is available at https://github.com/AWMurrayLab/FluctuationTest_GregLang. The basic LD model used by Lang *et al*. is equivalent to that available in the rsalvador package (using the function newton.LD), except without the correction for partial plating; the latter is only important when using the estimate of *m* to infer the mutation rate, not when comparing the fits of different models to the empirical cumulative distribution of the raw colony counts.

Plots of the empirical cumulative distributions of our data against the theoretical models showed that the Hamon and Ycart model was a visually good fit in all cases (see Fig. S9 and S11 for examples). To further assess goodness-of-fit of the Hamon and Ycart model, we performed Pearson’s chi-square test in R after binning the data and theoretical distribution such that the expected number of cultures in each bin of mutant counts was at least five^79^. To compare the goodness-of-fit of the Hamon and Ycart model to the basic LD model, we calculated the negative log-likelihood for each model and performed the likelihood ratio test. To compare the Hamon and Ycart model to the mixed LD-Poisson model, we simply compared the negative log-likelihoods (smaller values indicate a better fit); the likelihood ratio test was not applicable as these two models contain the same number of parameters. Note that although the Hamon and Ycart (or in some cases, LD-Poisson) models were often better fits than the basic LD model, we still used the basic LD model for statistical comparison of mutation rates between conditions, because an accurate method to account for partial plating has not yet been developed for the cases of post-plating mutations or differential fitness between mutants and parent strains^77^. Nevertheless, similar patterns in mutation rates were observed when using an older method of accounting for partial plating to derive µ_app_ from the Hamon and Ycart model^80^, indicating that the effects of PYO were robust to different mathematical methods of inferring mutation rates from fluctuation test data (Table S2).

### Characterization of antibiotic resistance phenotypes

We defined putative ciprofloxacin-resistant mutants as “enriched” by PYO in the fluctuation tests if colonies with a given morphology were at least 2x more numerous on the PYO-containing ciprofloxacin plate than the respective non-PYO-containing ciprofloxacin plate derived from the same 200 µL culture. These putative mutants could be either from the PYO pre-treated or non-PYO pre-treated branches of the fluctuation test. Putative mutants that were seemingly enriched by PYO were restreaked for purity on PYO-containing agar plates at the same ciprofloxacin concentration on which they were selected in the fluctuation test (0.5 µg/mL for *PA*, 8 µg/mL for *B. multivorans*). Putative mutants that were not enriched by PYO were restreaked on ciprofloxacin agar plates without PYO. Frozen stocks of each restreaked, visually pure isolate were prepared by inoculating cultures with single colonies in 5 mL of liquid LB, incubating to stationary phase, mixing 1:1 with 50% glycerol, and storing at -80°C.

The levels of ciprofloxacin resistance of selected isolates, as well as the parent strains, were assessed using a CFU recovery assay as follows. For each isolate, four 5 mL cultures in GMM (for *P. aeruginosa*) or GMM + AA (for *B. multivorans*) were inoculated directly from the frozen stock, to minimize the number of generations in which secondary mutations could be acquired. The cultures were grown to stationary phase overnight, then subcultured to an OD_500_ of 0.05 in 5 mL of fresh GMM (for *P. aeruginosa*) or GMM + AA (for *B. multivorans*), with or without 100 µM PYO. The new cultures were grown to mid log-phase, then serially diluted in GMM or GMM + AA (+/- 100 µM PYO as appropriate) and plated for CFUs (10 µL per dilution step) on 1) plain MH agar, 2) MH agar + ciprofloxacin, and 3) MH agar + ciprofloxacin + 100 µM PYO. The lowest plated dilution was 10^−1^, making the limit of detection approximately 1000 CFUs/mL.

### Identification of mutations by whole-genome sequencing

Genomic DNA was isolated from selected putative mutants and the parent strains using the DNeasy Blood & Tissue kit (Qiagen). Library preparation and 2×150 bp paired-end Illumina sequencing was performed by the Microbial Genome Sequencing Center (Pittsburgh, PA), with a minimum of 300 Mb sequencing output per sample (∼50x coverage). Forward and reverse sequencing reads were concatenated into a single file for each isolate and quality control was performed using Trimmomatic (version 0.39)^81^ with the following settings: LEADING:27 TRAILING:27 SLIDINGWINDOW:4:27 MINLEN:35. Mutations were then identified using breseq (version 0.34.1)^82^. The annotated reference genome for *P. aeruginosa* UCBPP-PA14 was obtained from BioProject accession number PRJNA38507. For *B. multivorans* AU42096, no reference genome was available from NCBI. Therefore, a genome scaffold was assembled from the paired-end sequencing data for the parent strain using SPAdes (version 3.14.0) with default parameters^83^. This scaffold was then used as the reference for breseq. Differences between the parent strain and isolates were identified using the gdtools utility that comes with breseq to compare the respective breseq outputs. All sequenced *P. aeruginosa* mutants were derived from the Δ*phz* strain except for CipR-33 and CipR-40, which were derived from the WT strain. In the case of *B. multivorans*, several dozen putative mutations were identified that were common to all three sequenced putative mutants. We assumed that these represented assembly errors in the parent strain genome scaffold, but even if they were genuine mutations, these would not account for the phenotypic differences between the isolates; therefore, Table S5 reports only mutations that were unique to each isolate. The genomic loci containing each putative mutation for the *B. multivorans* isolates were identified by retrieving the surrounding 200 bp from the parent genome scaffold and using the nucleotide BLAST tool on the MicroScope platform^84^ to find the closest match in the *B. multivorans* ATCC 17616 genome.

### Statistical analyses

All statistical analyses were performed using R^72^. Welch’s unpaired t-tests or one-way ANOVA with post hoc Tukey’s HSD test for multiple comparisons were used for tolerance assay data. Welch’s paired t-tests with Benjamini-Hochberg corrections for controlling false discovery rates were used for comparisons of apparent mutation rates across different conditions. Welch’s unpaired t-tests with Benjamini-Hochberg corrections for controlling false discovery rates were used for comparisons of CFU recovery on ciprofloxacin plates under different PYO treatments. For all antibiotic tolerance assays measured by CFUs, survival data were log_10_-transformed before statistical analyses.

