## Supplemental Text & Figures for "Bacterial defenses against a natural antibiotic promote collateral resilience to clinical antibiotics"

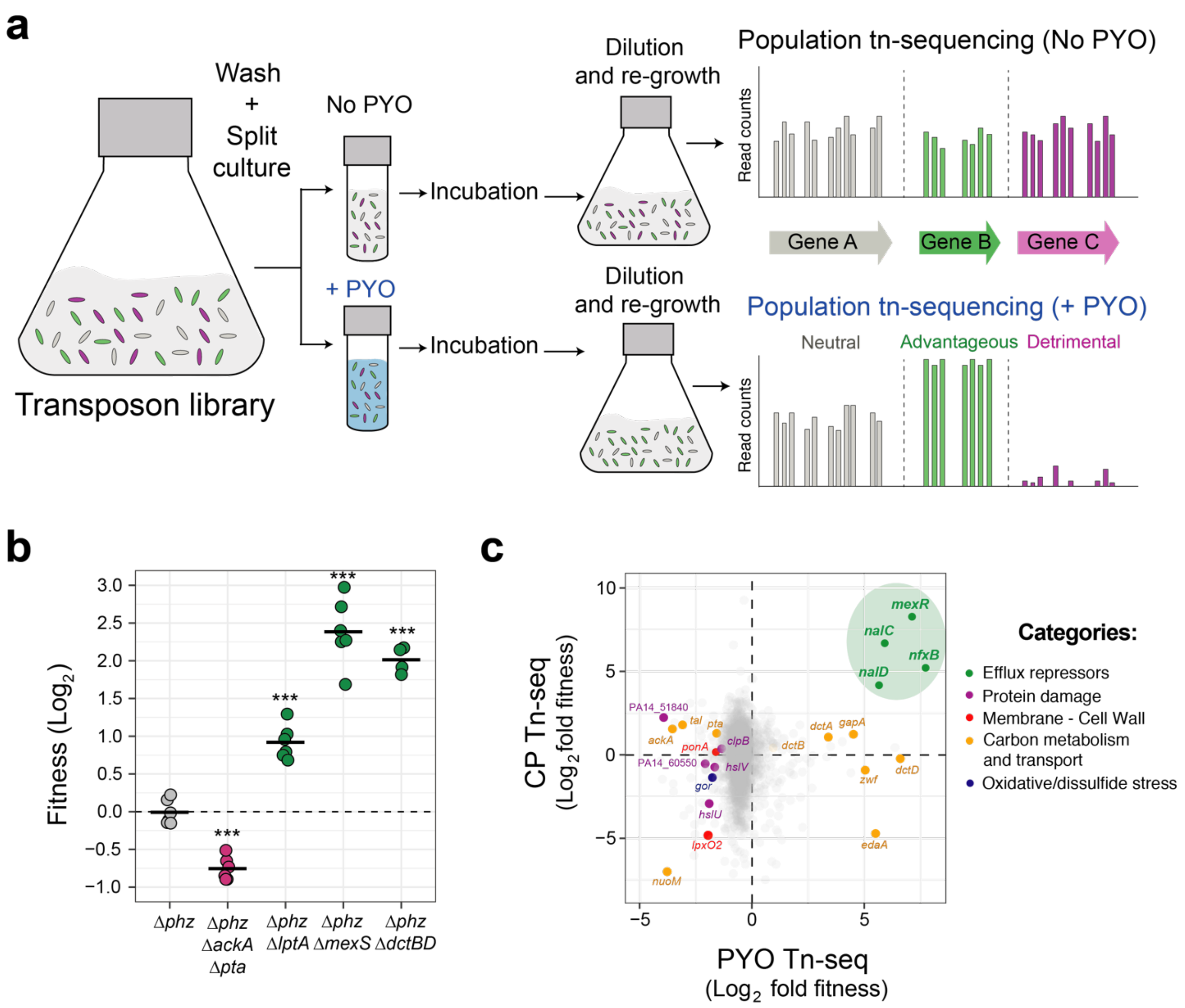


**Fig. S1. Tn-seq results and validations.** **A.** Tn-seq experimental design. Cells were incubated with and without PYO under nutrient starvation for maximum PYO toxicity[8] (see Methods for details). Bar graphs shown are hypothetical representations of the expected results for genes with different fitness effects and are not derived from the obtained data. **B.** Tn-seq validations. Chromosomal clean deletion mutants were exposed to PYO under carbon starvation, similar to the conditions used for the Tn-seq experiment. Survival of each strain was measured by colony forming units (CFUs) and compared to the survival of the parent Δ*phz* strain for fitness calculation (see Methods for details). Statistical significance was calculated using one-way ANOVA with

Tukey's HSD multiple-comparison test, with asterisks showing significant differences relative to Δ*phz* (*** $p < 0.001$). Data points represent independent biological replicates and black horizontal lines mark the mean fitness for each strain. **C.** Complete correlation analysis between PYO Tn-seq (this study) and ciprofloxacin (CP) Tn-seq[14]. Genes shown in Fig. 1A that were also present the CP Tn-seq dataset are highlighted in the plot. Efflux repressors are highlighted within the green circle. For full comparison of fitness values for both datasets, see Table S1.

<u>Additional discussion about Tn-seq data (Figs. 1A, S1B and S1C)</u>: The lack of any correlation for genes within the "Carbon metabolism and transport" category is expected since each Tn-seq experiment was performed in a different growth medium (LB for CP Tn-seq vs. succinate minimal medium for PYO Tn-seq; see Methods). In our PYO Tn-seq, disruptions in some of these genes (such as the *dctBD* two-component system that senses and transports succinate[85]) increased fitness, while disruptions in others (such as *nuoM*, a component of the electron transport chain[86]) decreased fitness. These complex results likely reflect the fact that in the presence of ROS-generating toxins like PYO, the cell must balance conflicting priorities: it needs to maintain proton-motive force to pump the toxins out, and have NADH available to power reductases involved in repair of oxidative damage, but at the same time it might benefit from limiting flux through the electron transport chain to decrease the potential for ROS generation. Our results hint that tuning cellular metabolism to an optimal compromise is carbon source dependent, which has also previously been suggested in a study on susceptibility to aminoglycosides[87]; the details of how such a compromise is regulated are worthy of future investigation. As for the other categories of genes that were implicated in tolerance to PYO, accumulation of PYO inside the cells might damage proteins through oxidative side reactions, which would explain why mutants in some of the protein damage-repair systems showed decreased fitness. Glutathione is also essential for protection against oxidative stress

damage[88] and disruptions in genes involved in its biosynthesis (*gshA*, *gshB*) or reduction (*gor*) decreased fitness (Fig. 1A). Finally, for genes related to cell wall/membrane synthesis, it is possible that the transposon insertions, by altering permeability, caused either an increase or decrease of how much PYO went into the cell. An increased influx of PYO would exacerbate cell damage, while decreased permeability would reduce PYO influx and alleviate toxicity.

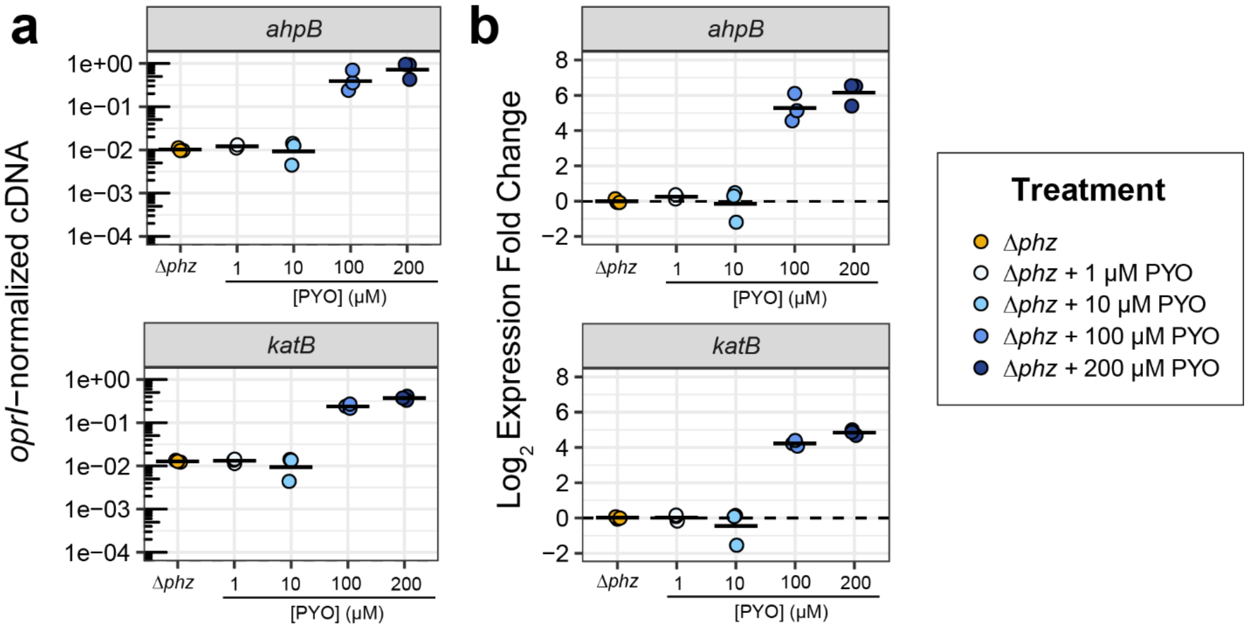


**Fig. S2. Effects of different concentrations of PYO on the expression of the *P. aeruginosa* oxidative stress response genes *ahpB* and *katB*. A.** Normalized cDNA levels measured by qRT-PCR. cDNA measurements were normalized by levels of the housekeeping gene *oprI* (see Methods). **B.** Fold change in expression upon PYO treatment, relative to the measurements in untreated Δ*phz*. *ahpB*: alkyl hydroperoxide reductase B; *katB*: catalase B. Black horizontal lines mark the mean value for independent biological cultures (n = 3).

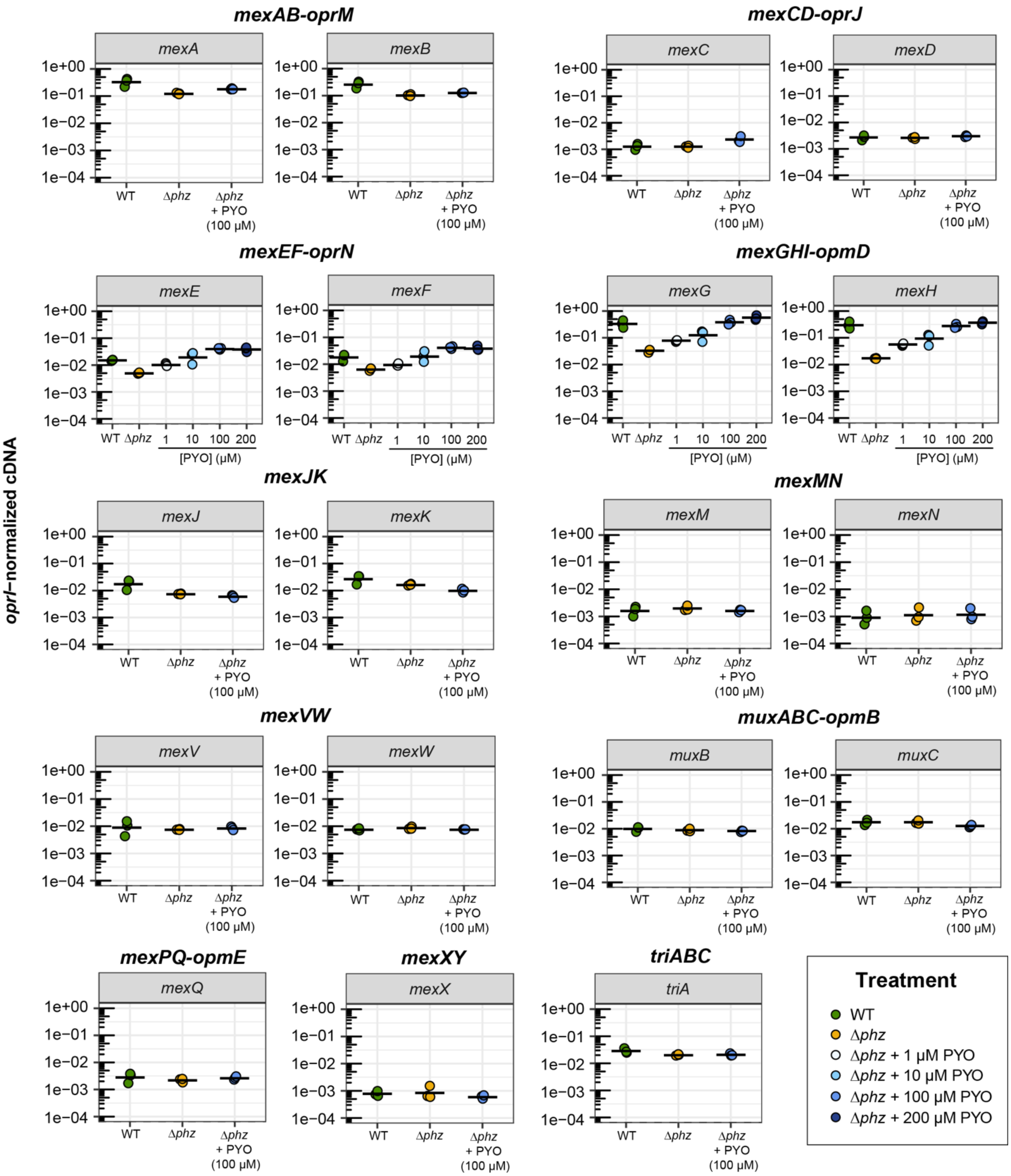


**Fig. S3. Effects of PYO on the expression of *P. aeruginosa* RND efflux systems (normalized cDNA levels)**. The normalized cDNA levels for genes within operons coding for the 11 main RND efflux systems in *P. aeruginosa* are shown. cDNA levels for each gene were measured by qRT-PCR during early stationary phase and normalized by the levels of the housekeeping gene *oprI* (see Methods). This dataset was used to make the heatmap presented in Fig. 2B. Black horizontal lines mark the mean value for independent biological cultures (n = 3).

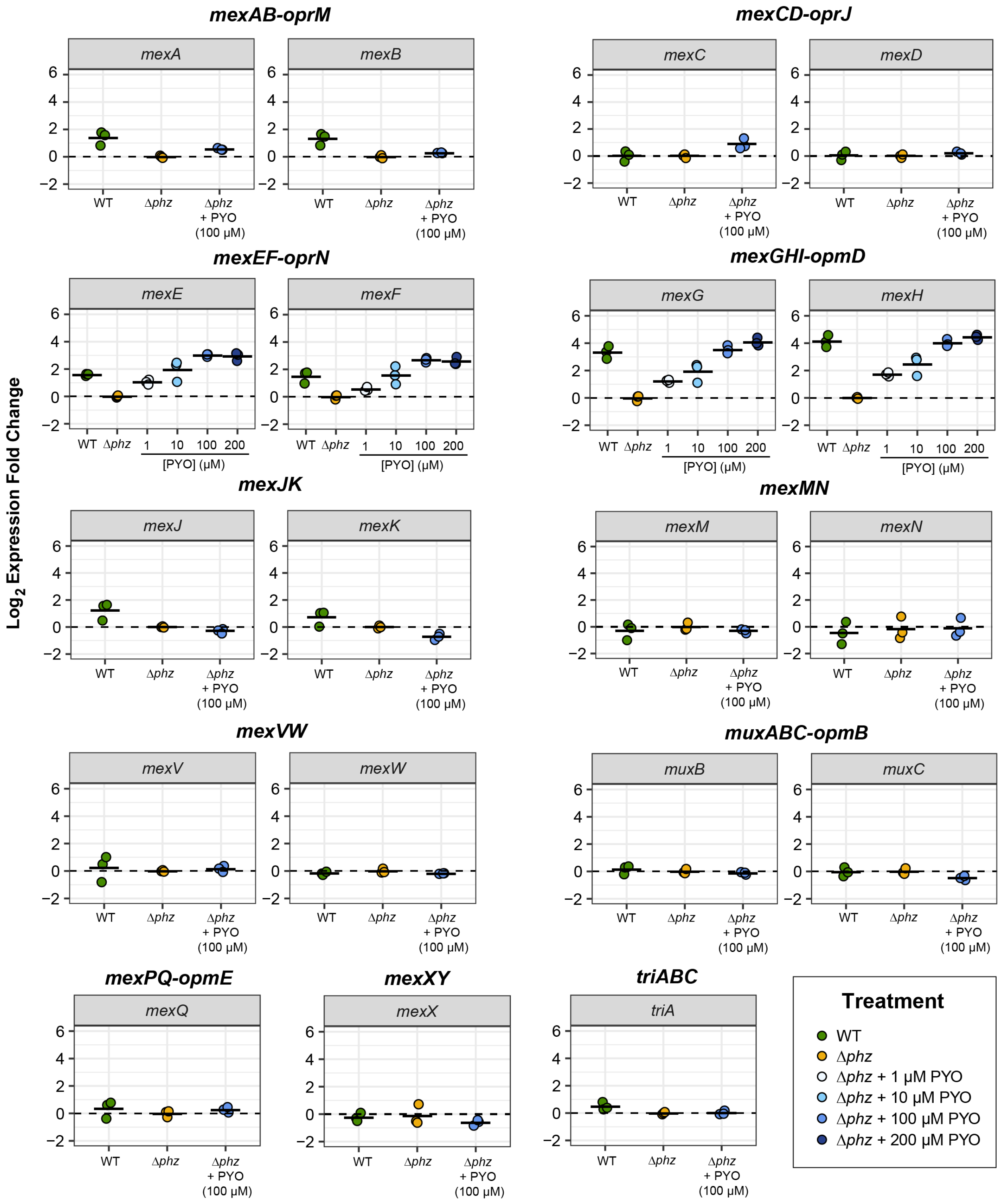


**Fig. S4. Effects of PYO on the expression of *P. aeruginosa* RND efflux systems (fold change).** The PYO-induced changes in expression for genes within operons coding for the 11 main RND efflux systems in *PA* are shown. These plots are derived from the normalized cDNA dataset shown in Fig S3. Here, the values for Δ*phz* were used as the basis for calculation of changes of expression (shown as $\log_2$ fold change). Black horizontal lines mark the mean value for independent biological cultures (n = 3).

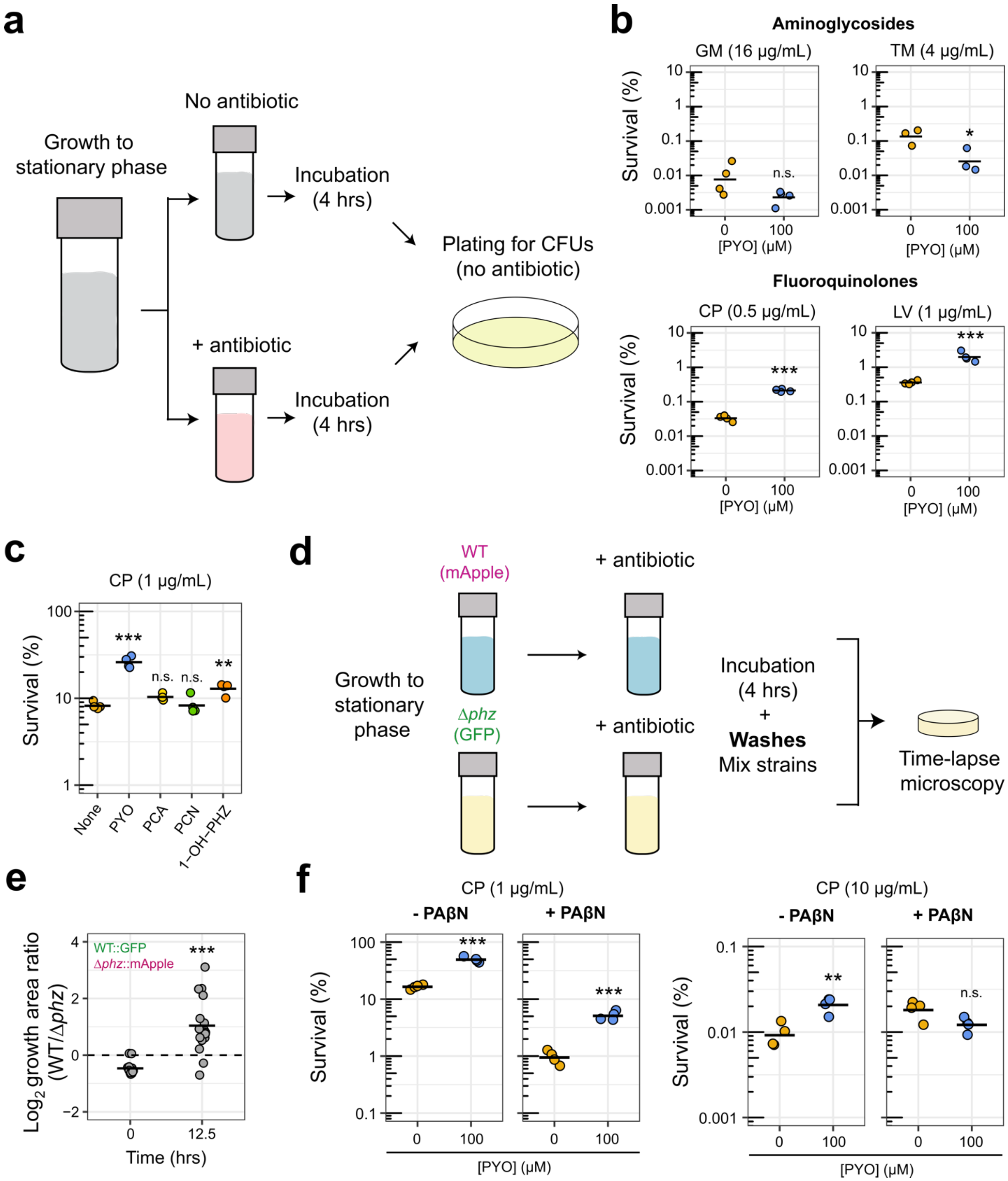


**Fig. S5. Effects of PYO on *P. aeruginosa* tolerance to different antibiotics**. **A.** Experimental design for survival assay to measure tolerance to clinical antibiotics. In conditions with exogenous PYO, the PYO was added when cultures were inoculated. PYO itself was not lethal under these experimental conditions. **B.** Tolerance levels of Δ*phz* cells harvested in log phase, following growth in the presence or absence of PYO (100 μM), to different aminoglycosides and fluoroquinolones (GM = gentamicin, TM = tobramycin, CP = ciprofloxacin, and LV = levofloxacin). Data points represent independent biological replicates (n = 4 for all except tobramycin, for which n = 3). Stationary phase tolerance experiments are not shown for the aminoglycosides (gentamicin and tobramycin), as treatment with tobramycin in stationary phase under our conditions at this clinically-relevant concentration[25] did not result in cell death, regardless of the presence of PYO. **C.** Effect on tolerance to CP caused by the presence of the four main phenazines produced by *P. aeruginosa* (PYO = pyocyanin, PCA = phenazine-1-carboxylic acid, PCN = phenazine-1-carboxamide and 1-OH-PHZ = 1-hydroxyphenazine). Cultures were in stationary phase when exposed to CP, and tolerance assays were performed as shown in panel A (see Methods) (n = 4). These experiments were performed using a Δ*phz** mutant that can neither produce nor modify any of the phenazines. **D.** Experimental design for time-lapse microscopy experiments, in which cells were grown on agarose pads after exposure to CP (10 μg/mL). The strain/fluorescent protein examples shown (i.e. WT::mApple, Δ*phz*::GFP) are the ones used in the images of Fig. 2D and movie S1. **E.** Quantification of microscopy data as done in Fig. 2E, but for the experiment with swapped fluorescent proteins. **F.** Effects of the efflux inhibitor PAβN on tolerance levels to CP of Δ*phz* cells grown in the presence or absence of PYO (100 μM). Cultures were treated with low (left) and high (right) CP concentrations. Experiments for all the conditions were done in parallel (see Methods for full protocol) (n = 4). Statistics: B, E, F – Welch's unpaired t-tests. C – One-way ANOVA with Tukey's HSD multiple-comparison test, with asterisks showing the statistical significance of comparisons with the untreated (no phenazines) (* $p < 0.05$, ** p <

0.01, *** $p < 0.001$, n.s. $p > 0.05$). In panels B, C, E and F, black horizontal lines mark the mean value for independent biological cultures.

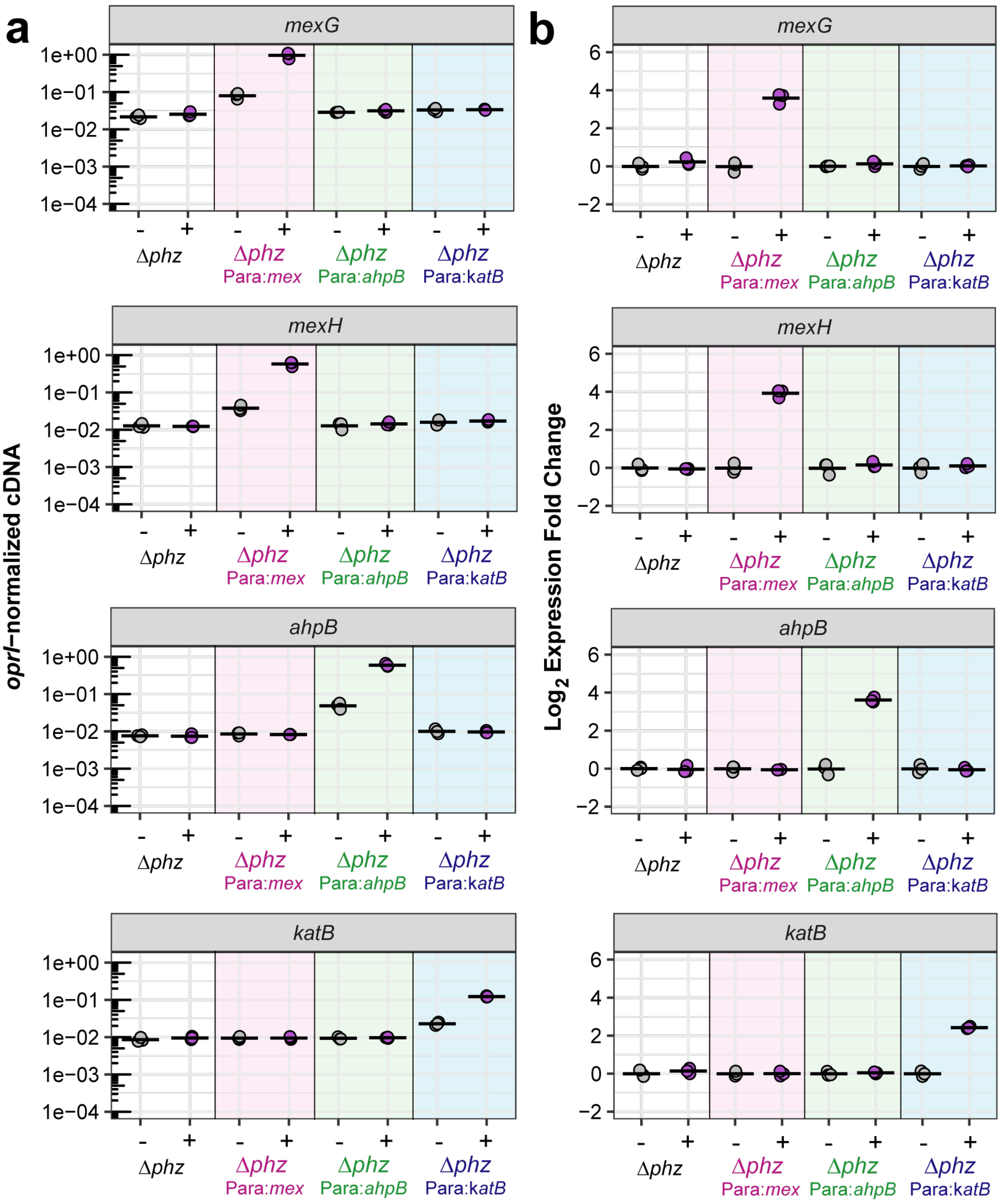


**Fig. S6. Artificial induction of *mexGHI-opmD*, *ahpB* and *katB*. A.** Normalized cDNA levels measured by qRT-PCR. cDNA levels were normalized by the housekeeping gene *oprI*. **B.** Fold change in expression upon arabinose induction. This dataset can be compared to the PYO-mediated induction of the same genes as shown in Figs. 2A, S2 and S3. The four strains shown are: 1) the parent Δ*phz* (white background), 2) Δ*phz* $P_{ara}$:*mexGHI-opmD* (magenta background), 3)

Δ*phz* $P_{ara}$:*ahpB* (green background) and 4) Δ*phz* $P_{ara}$:*katB* (blue background). +/- represent addition or not of 20 mM arabinose to the cultures for the artificial induction of expression. For additional experimental details and strain information, see Methods and Table S4. Black horizontal lines mark the mean value for independent biological cultures (n = 3).

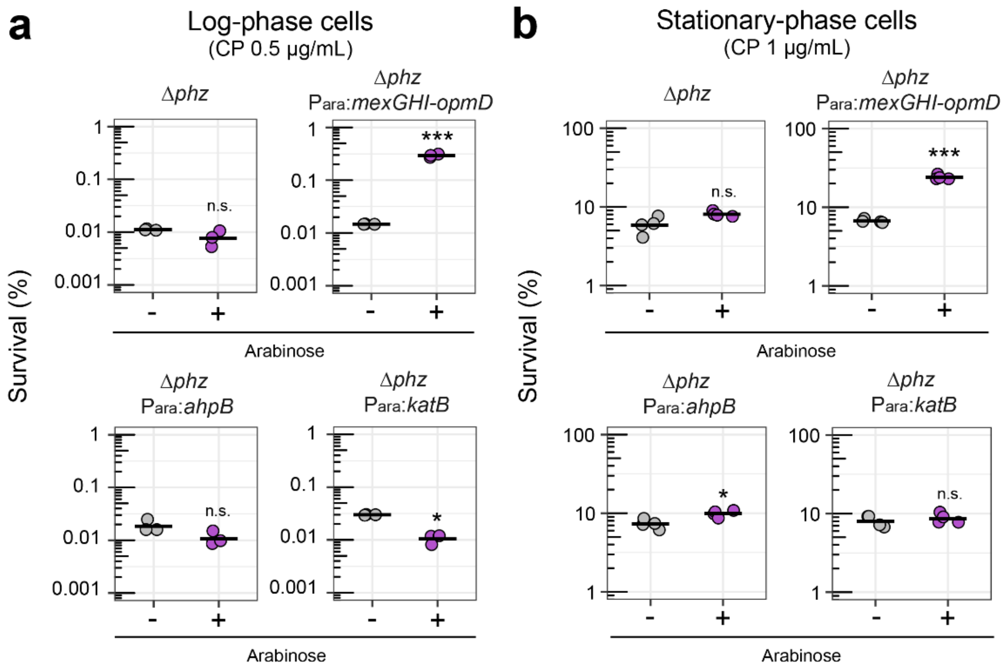


**Fig. S7. Effect of artificial induction of PYO-induced genes on tolerance to ciprofloxacin.** Survival relative to the no-antibiotic control is shown for the parent Δ*phz* strain and the three arabinose-inducible strains (in which the PYO-inducible genes *mexGHI-opmD, ahpB,* or *katB* are under control of an arabinose-inducible promoter) grown in the presence or absence of 20 mM arabinose, without exposure to PYO. The tolerance experiments were performed for cultures in both log phase (A, n = 3) and stationary phase (B, n = 4). In B, the experiment for *mexGHI-opmD* is the same as in Fig. 2G, but is also shown here for ease of comparison. Statistics: Welch's unpaired t-tests (* $p < 0.05$, ** $p < 0.01$, *** $p < 0.001$, n.s. $p > 0.05$). Black horizontal lines mark the mean value for independent biological cultures.

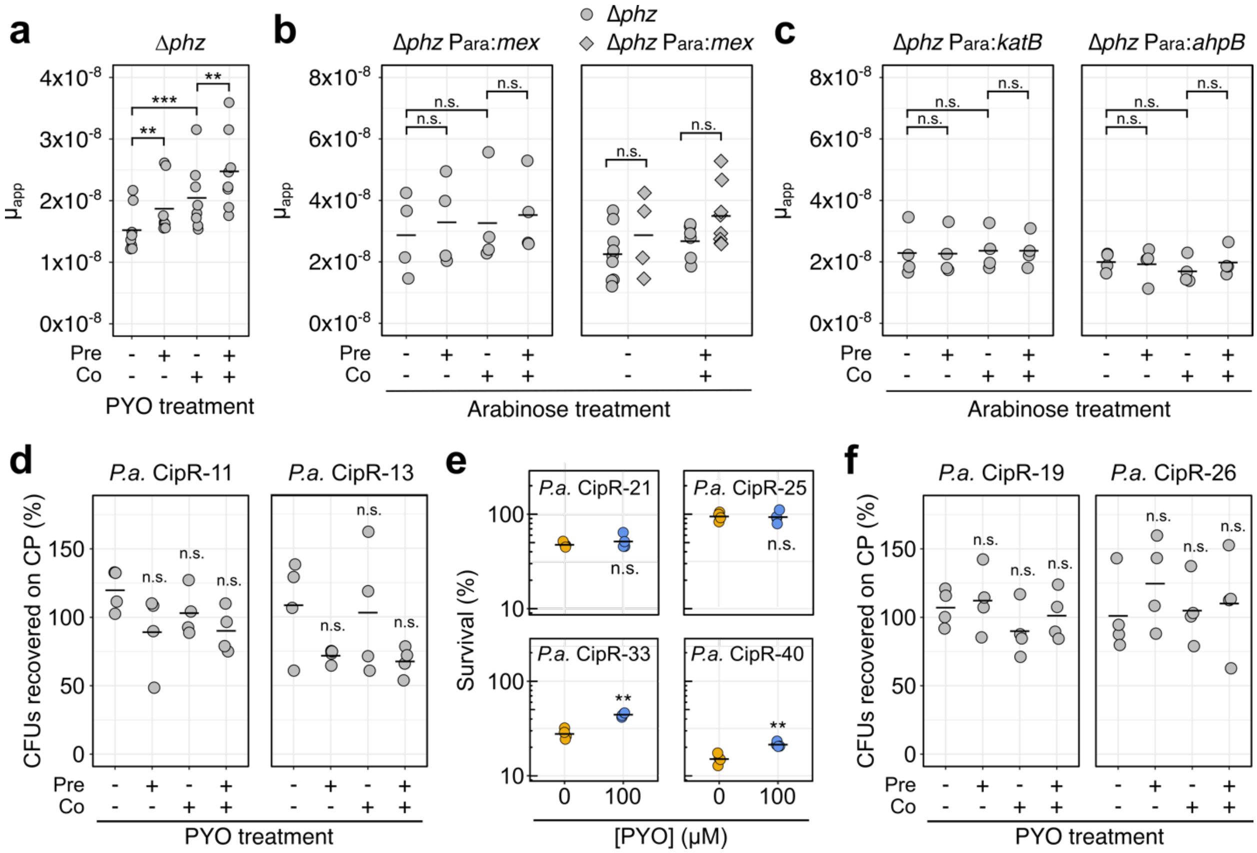


**Fig. S8. Effect of PYO or PYO-induced genes on apparent mutation rates, resistance phenotypes, and antibiotic tolerance of partially-resistant mutants. A.** Apparent mutation rates of stationary-phase Δ*phz* grown in liquid minimal medium and plated onto MH agar containing ciprofloxacin (CP, 0.5 µg/mL), with or without pre- and/or co-exposure to 100 µM PYO relative to the antibiotic selection step. Each data point represents 44 parallel cultures from a single biological replicate (n = 8). **B-C.** Apparent mutation rates of log-phase cells grown in glucose minimal medium and plated onto MH agar containing CP (0.5 µg/mL), with or without pre- and/or co-exposure to 20 mM arabinose relative to the antibiotic selection step. Data are shown for Δ*phz* $P_{ara}$:*mexGHI-opmD* alone (B, left, n = 4) and in comparison to Δ*phz* (B, right), or Δ*phz* $P_{ara}$:*katB* and Δ*phz* $P_{ara}$:*ahpB* (C, n = 4). Each data point represents 44 parallel cultures from a single biological replicate. **D.** The percentage of CFUs recovered on CP (0.5 µg/mL) either with or without 100 µM PYO in the agar, for log-phase cultures of representative resistant mutants that were not enriched by exposure to PYO in the fluctuation tests. The mutants were pre-grown with or without 100 µM PYO in glucose minimal medium before plating. Percentage recovery was calculated relative to total CFUs counted on non-selective plates (n = 4). **E.** Tolerance to CP (1 µg/mL) of partially-resistant mutants grown in glucose minimal medium to stationary phase with or without 100 µM PYO (n = 4). Experiments were performed as shown in Fig. S5A. **F.** The percentage of CFUs recovered on CP (0.5 µg/mL) for log-phase cultures of representative resistant mutants that were enriched by exposure to PYO in the fluctuation tests (n = 4). Experiments were performed in the same way as in panel D. Statistics: A – Welch's paired t-tests with Benjamini-Hochberg correction for controlling false discovery rate; B-F – Welch's unpaired t-tests (* $p < 0.05$, ** $p < 0.01$, *** $p < 0.001$, n.s. $p > 0.05$). Unless indicated otherwise with brackets, statistical significance is shown for the comparison with the untreated (no PYO) condition. Data points represent independent biological cultures, with horizontal black lines marking the mean value for each condition.

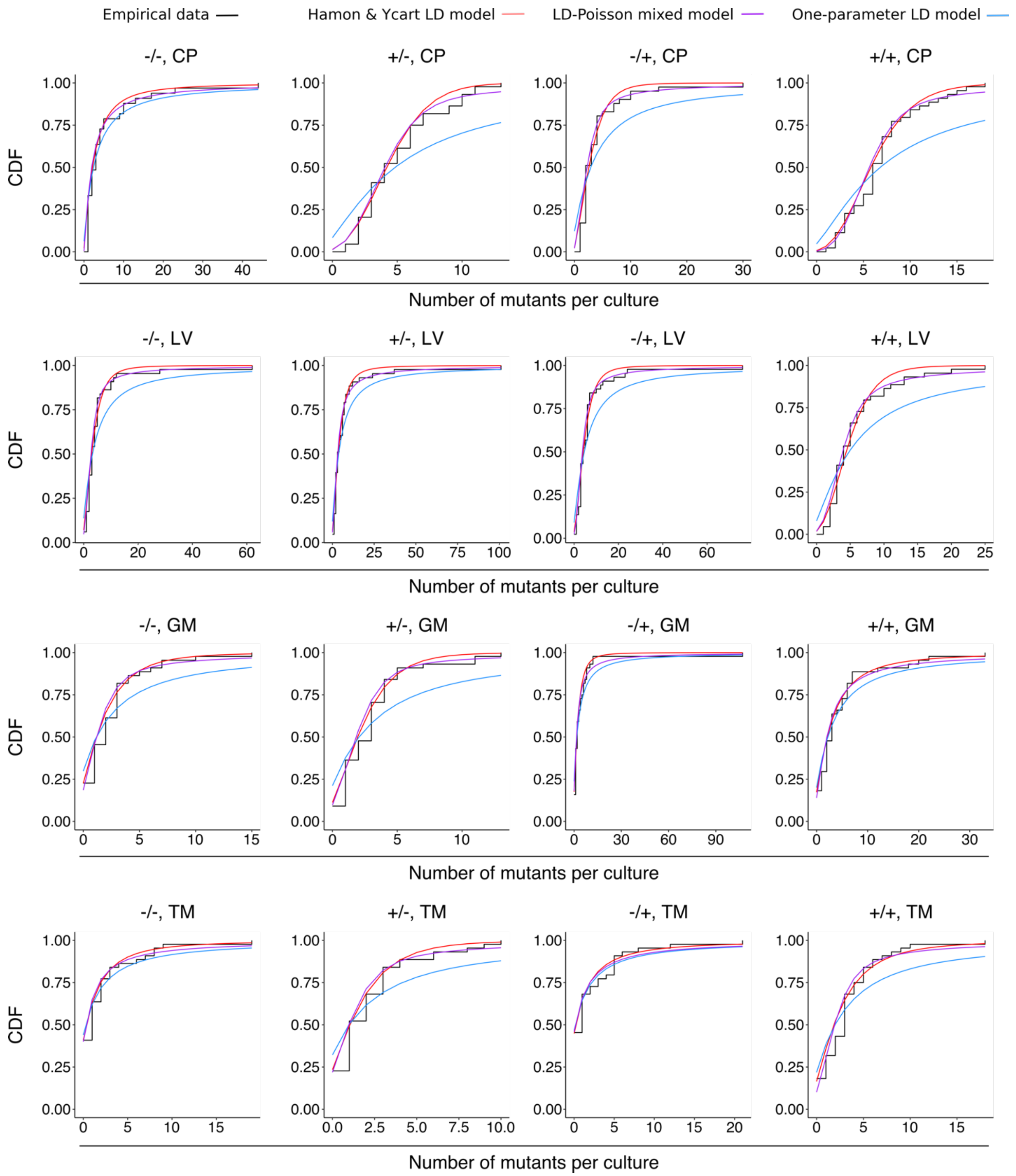


**Fig. S9. Goodness-of-fit of different mathematical models for *P. aeruginosa* Δ*phz* fluctuation test data.** Data are plotted for different combinations of PYO in liquid (pre-treatment) / PYO in agar (co-exposure to antibiotic selection). The empirical cumulative distribution functions of the data (black) are plotted against 1) a variation of the Luria-Delbrück model fit with two parameters, *m* (the expected number of mutations per culture) and *w* (the relative fitness of mutant cells vs. WT), as implemented by Hamon & Ycart[32]; 2) a mixed Luria-Delbrück and Poisson distribution fit with two parameters, *m* and *d* (the number of generations that occur post-plating), allowing for the possibility of post-plating mutations, as implemented by Lang *et al.*[33]; 3) the basic Luria-Delbrück distribution model (blue) fit only with *m*, as implemented by Lang *et al.*[33] In each condition, the plotted data represent the biological replicate with the lowest chi-square goodness-of-fit *p*-value (i.e. least-good fit) for the Hamon & Ycart model.

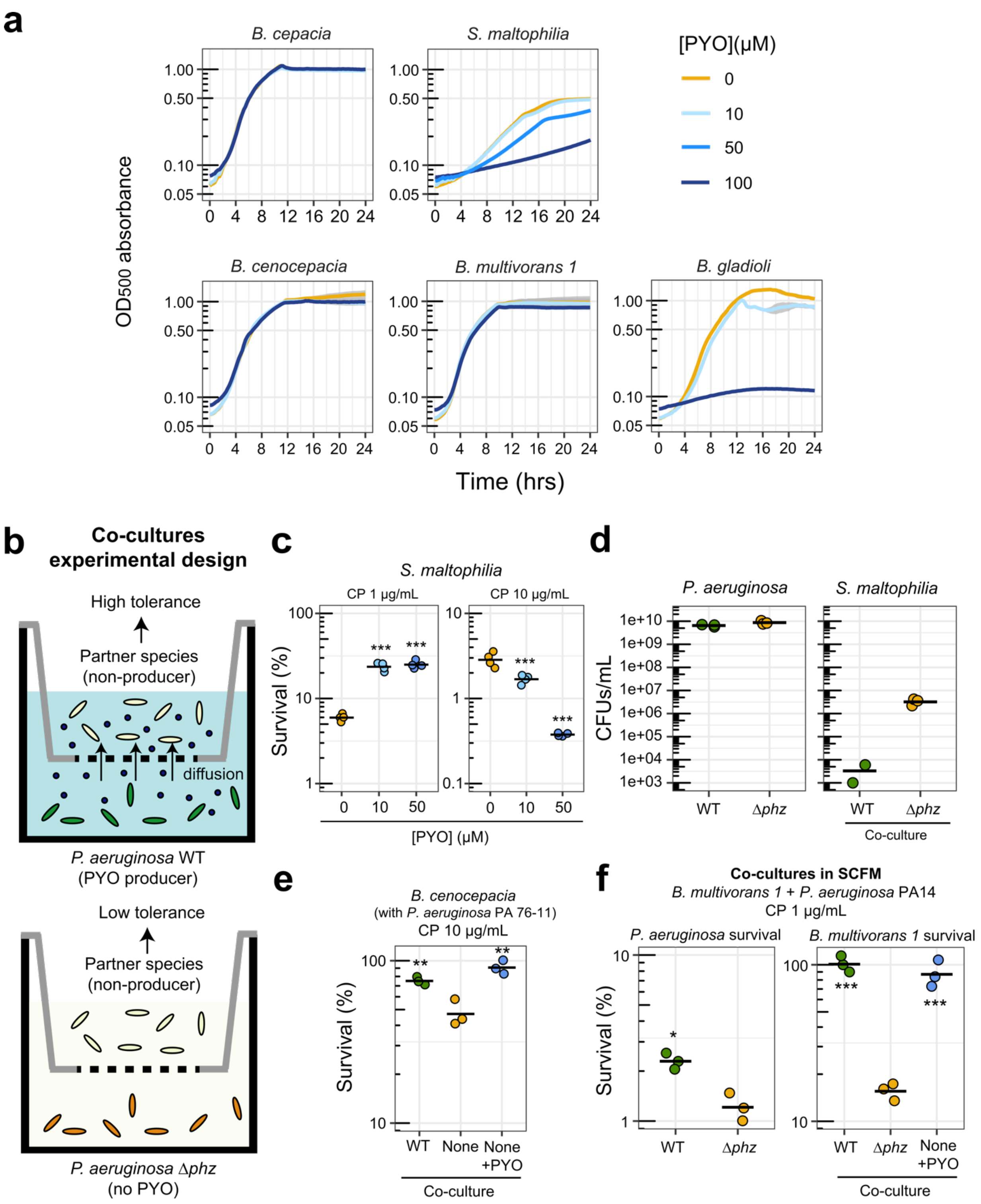


**Fig. S10. Effects of PYO on growth and antibiotic tolerance of non-phenazine producing opportunistic pathogens**. **A.** Growth of several strains in the presence of different concentrations of PYO. Plotted lines represent averages of four to six replicates and shaded areas in gray represent the standard deviation. *Burkholderia multivorans 1* = *B. multivorans* AU42096. For complete information on strains, see Table S4. **B.** Schematic depicting the experimental design for co-culture antibiotic tolerance assays. **C.** Tolerance of *S. maltophilia* to different concentrations of ciprofloxacin (CP; 1 or 10 μg/mL) after growth in the presence of different concentrations of PYO (0, 10 or 50 μM) (n = 4). The panel on the right is the same as in Fig. 4C, but is also shown here for ease of comparison. **D.** CFUs recovered from co-cultures of *P. aeruginosa* (PA14 WT and Δ*phz*) and *S. maltophilia* (n = 3). **E**. Tolerance levels to CP of *B. cenocepacia* when grown in co-culture with a PYO-producing clinical *P. aeruginosa* strain (PA 76-11) or when grown alone with or without PYO (100 μM) added exogenously (n = 3). **F**. Tolerance to CP of *P. aeruginosa* (WT and Δ*phz* PA14 grown in co-cultures with *B. multivorans 1*, left) and *B. multivorans 1* (grown in co-cultures with PA14 WT or Δ*phz* or alone with 100 μM PYO added exogenously, right) (n = 3). This experiment was performed in SCFM. Statistics: C, E, F (right) - One-way ANOVA with Tukey's HSD multiple-comparison test, with asterisks showing the statistical significance of comparisons with the untreated sample (no PYO). F (left) - Welch's unpaired t-test (* $p < 0.05$, ** $p < 0.01$, *** $p < 0.001$, n.s. $p > 0.05$). In panels C-F, individual data points represent independent biological replicates and black horizontal lines mark the mean value for each condition.

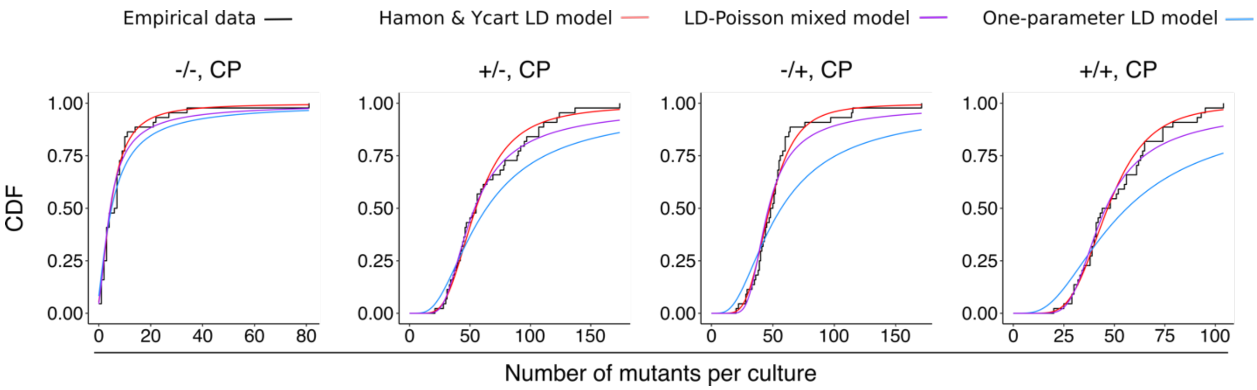


**Fig. S11. Goodness-of-fit of different models for *B. multivorans* fluctuation test data.** Data are plotted for different combinations of PYO in liquid (pre-treatment) / PYO in agar (co-exposure to antibiotic selection). The empirical cumulative distribution functions of the data (black) are plotted against 1) a variation of the Luria-Delbrück model fit with two parameters, *m* (the expected number of mutations per culture) and *w* (the relative fitness of mutant cells vs. WT), as implemented by Hamon & Ycart[32]; 2) a mixed Luria-Delbrück and Poisson distribution fit with two parameters, *m* and *d* (the number of generations that occur post-plating), allowing for the possibility of post-plating mutations, as implemented by Lang *et al.*[33]; 3) the basic Luria-Delbrück distribution model (blue) fit only with *m*, as implemented by Lang *et al*.[33] In each condition, the plotted data represent the biological replicate with the lowest chi-square goodness-of-fit *p*-value (i.e. least-good fit) for the Hamon & Ycart model.

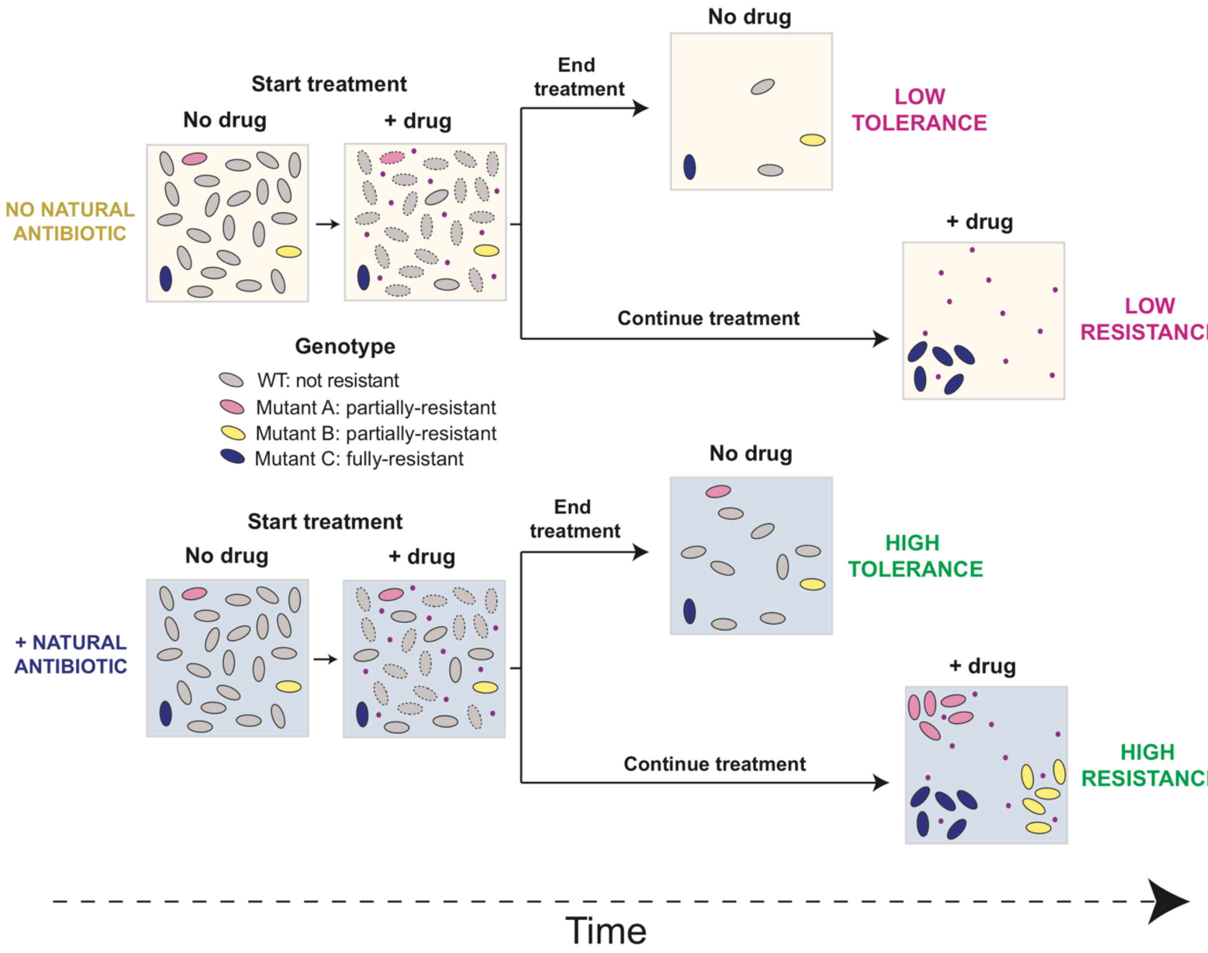


**Fig. S12. Proposed model for how exposure to natural antibiotics increases bacterial tolerance and resistance to clinical drugs**. In the first scenario (tolerance), cells are exposed to the clinical drug (pink dots) for a short period of time. Surviving cells will eventually re-start growth after the drug is removed. The presence of the natural antibiotic (bottom) increases tolerance of both WT and partially-resistant mutants. In the second scenario (resistance), cells are constantly exposed to the drug for an extended period of time, and only mutants are maintained in the population. The presence of the natural antibiotic (bottom) allows partially-resistant mutants to grow under drug selection, preserving a greater range of genetic diversity in the population.
